# Structural basis for template switching by a group II intron-encoded non-LTR-retroelement reverse transcriptase

**DOI:** 10.1101/2021.05.13.443781

**Authors:** Alfred M. Lentzsch, Jennifer L. Stamos, Jun Yao, Rick Russell, Alan M. Lambowitz

## Abstract

Reverse transcriptases (RTs) can template switch during cDNA synthesis, enabling them to join discontinuous nucleic acid sequences. Template switching plays crucial roles in retroviral replication and recombination, is used for adapter addition in RNA-seq, and may contribute to retroelement fitness by enabling continuous cDNA synthesis on damaged templates. Here, we determined an X-ray crystal structure of a template-switching complex of a group II intron RT bound simultaneously to an acceptor RNA and donor RNA template/DNA heteroduplex with a 1-nt 3’-DNA overhang. The latter mimics a completed cDNA after non-templated addition (NTA) of a nucleotide complementary to the 3’ nucleotide of the acceptor as required for efficient template switching. The structure showed that the 3’ end of the acceptor RNA binds in a pocket formed by an N-terminal extension (NTE) present in non-long-terminal-repeat (LTR)-retroelement RTs and the RT fingertips loop, with the 3’ nucleotide of the acceptor base paired to the 1-nt 3’-DNA overhang and its penultimate nucleotide base paired to the incoming dNTP at the RT active site. Analysis of structure-guided mutations identified amino acids that contribute to acceptor RNA binding and a phenylalanine near the RT active site that mediates NTA. Mutation of the latter residue decreased multiple sequential template switches in RNA-seq. Our results provide new insights into the mechanisms of template switching and NTA by RTs, suggest how these reactions could be improved for RNA-seq, and reveal common structural features for template switching by non-LTR-retroelement RTs and viral RNA-dependent RNA polymerases.

## Introduction

Reverse transcriptases (RTs)^3^ are a large enzyme family, members of which function in the replication of human pathogens, such as retroviruses and hepatitis B virus, have played pivotal roles in the evolution of life on Earth, and are widely used for biotechnological applications, such as RT-PCR and RNA-seq (1–4). Although most studies have focused on retroviral enzymes and other RTs found in eukaryotes, RTs are thought to have originated in bacteria, likely from an RNA-dependent RNA polymerase (RdRP), and large numbers of RTs persist there, either encoded by retrotransposons called mobile group II introns or as free-standing genomically encoded enzymes (5–9). The latter are related to group II intron RTs but have acquired cellular functions, including phage/host tropism switching, acquisition of RNA spacers in CRISPR-Cas systems, and contributions to multiple types of phage defense mechanisms (10–13). These bacterial enzymes provide an evolutionary perspective on the biochemical properties and biological functions of RTs, which are much wider than suggested by a narrow focus on retroviral RTs.

Template switching is an important but as yet poorly understood biochemical activity of RTs and RdRPs. Studies of template switching by retroviral RTs focused initially on its role in retroviral replication and recombination and more recently on its use for adapter addition in RNA-seq (1,14–16). Template switching by retroviral RTs is dependent upon base pairing between the donor and acceptor nucleic acids and falls into two mechanistically distinct categories: strand transfers, which require long base-pairing interactions between the donor and acceptor nucleic acids, such as those involving long-terminal repeat (LTR) sequences during retroviral replication, and end-to-end clamping, which typically requires only two to four base pairs and is dependent upon non-templated nucleotide addition (NTA) to a completed cDNA (17–20). Retroviral RT clamping is employed in a widely used high-throughput RNA sequencing method called SMART-seq for 5’ RNA-seq adapter by template switching from the 5’ end of an RNA template to a synthetic adapter oligonucleotide whose 3’ end contains nucleotide residues complementary to those added by NTA to the 3’ end of a completed cDNA (14–16).

Group II intron and other bacterial RTs belong to a large subgroup of RTs that are encoded by non-LTR-retroelements and includes mitochondrial retroplasmid, human LINE-1 element, insect R2 element, and other eukaryotic non-LTR-retrotransposon RTs (2, 21). These non-LTR-retroelement RTs are homologous to retroviral RTs but have distinctive structural features, including a functionally important N-terminal extension (NTE) and two insertions, denoted RT2a and RT3a, between conserved RT sequence blocks in the RT domain (21, 22). An X-ray crystal structure of a full-length group II intron RT (the thermostable *Geobacillus stearothermophilus* GsI-IIC RT, sold commercially as TGIRT-III) bound to template-primer substrate and an incoming dNTP showed that these insertions contribute multiple additional contacts with RNA templates and more constrained binding sites for the templating RNA base, 3’ end of the DNA primer, and incoming dNTP that could contribute to the relatively high fidelity and processivity of group II intron RTs (23, 24). Structural cognates of all three of these distinctive regions are also found in RdRPs but are not present and presumed to have been lost from retroviral RTs, which evolved to evade host defenses by introducing frequent mutational variations and rapidly propagating beneficial ones by falling off one template and using the bound cDNA as a primer to reinitiate on another (24–26).

Studies in our laboratory aided by the crystal structure described above have focused on the template-switching activity of GsI-IIC RT (27). In a previous biochemical study, we used a series of synthetic acceptor RNAs and donor RNA template/DNA heteroduplexes representing the 5’ end of an RNA template annealed to a completed cDNA to establish a kinetic framework for the template-switching and related NTA reactions of this enzyme (27). These studies showed that template switching is most efficient from a donor RNA/DNA heteroduplex duplex with a 1-nt 3’-DNA overhang complementary to the 3’ nucleotide of the acceptor RNA, indicating a requirement for a single NTA to the 3’ end of the completed cDNA, and that the single base pair between the 1-nt 3’-DNA overhang and the 3’ nucleotide of the acceptor confers remarkably high specificity (97.5-99.7% precise junctions depending upon the base pair), as determined by high-throughput sequencing of the template-switching junctions (27).

Here, we determined a crystal structure of GsI-IIC RT in the act of template switching, to our knowledge the first such structure for any polymerase. The structure showed that the 3’ end of the acceptor RNA template binds in a pocket that is formed by the NTE and fingertips loop and is ordinarily occupied by upstream regions of a continuous RNA template. The structure provides new mechanistic insights, including how a single initial base-pairing interaction within the template-switching pocket could lead to highly specific template-switching junctions, and suggests how the template-switching and NTA reactions might be improved for RNA-seq applications. A similar template-switching pocket composed of a homologous NTE and fingertips loop is conserved in non-LTR-retroelement RTs and viral RdRPs, but only partially conserved in retroviral RTs, which lack the NTE and exhibit more promiscuous template-switching behavior.

## Results

### Structure of a group II intron RT template-switching complex

We determined a crystal structure of full-length *E. coli*-expressed GsI-IIC RT in complex with a 5-nt acceptor RNA (5′ rUrUrUrUrG), a donor duplex consisting of a 10-nt RNA template strand annealed to a complementary 11-nt DNA primer strand leaving a 1-nt 3’ dideoxy C overhang, and dATP (Fig. 1, *A* and *B* and Table S1). The donor duplex represents the 5’ end of an RNA template annealed to a completed cDNA after NTA of a nucleotide complementary to the 3’ nucleotide of the acceptor RNA but unable to initiate cDNA synthesis because of the 3’ dideoxy. Nucleotide residues in the structure are designated as belonging to the primer (P) or template (T) strands and are numbered positively or negatively from the dNTP-binding site at position -1, with T-1 (a U residue) corresponding to the templating RNA base for the incoming dATP (Fig. 1*B*).

**Figure 1.**
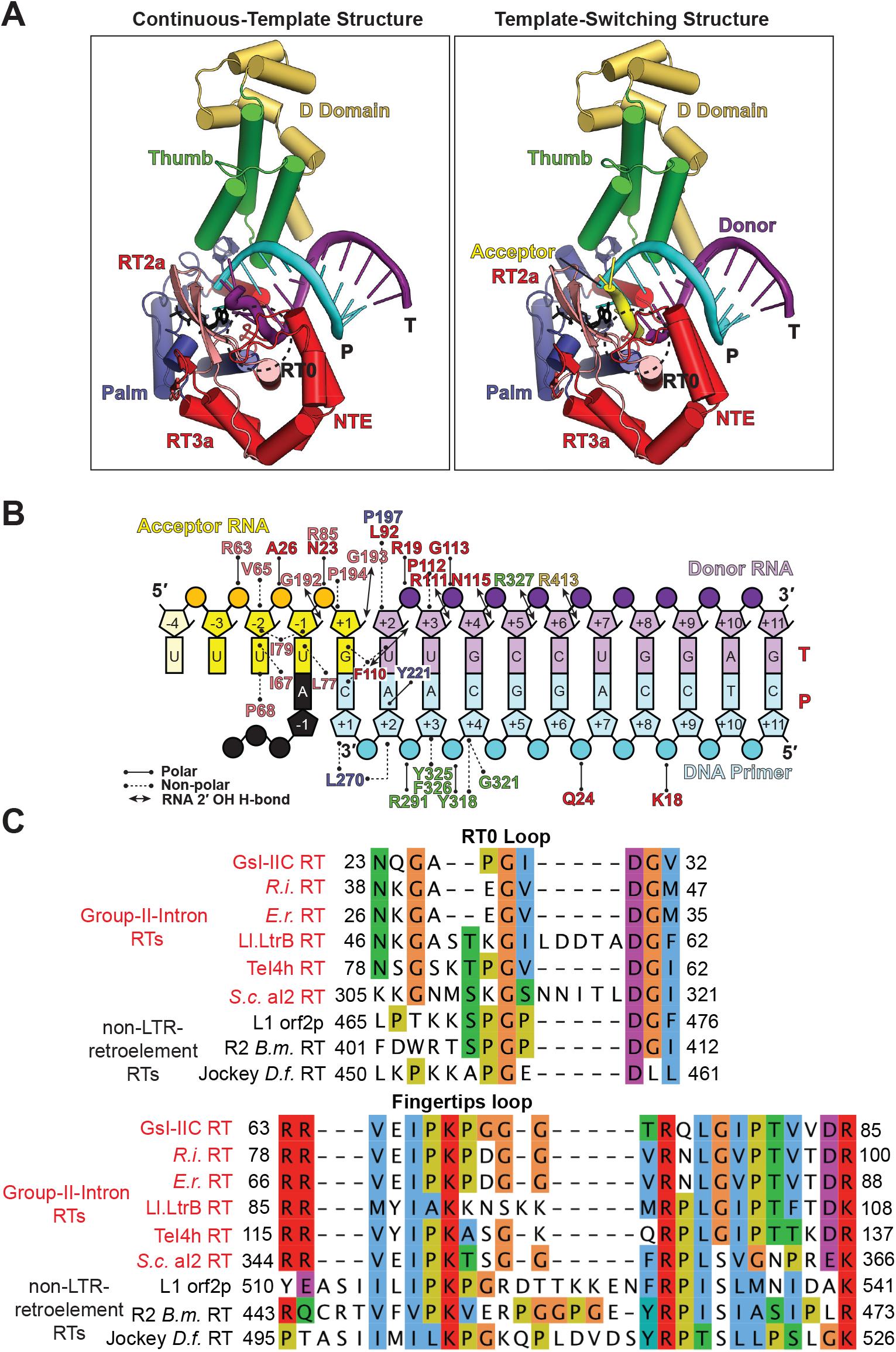
Structure of a group II intron RT template-switching complex. *A,* structure of GsI-IIC RT poised for template switching from a donor RNA template/DNA primer heteroduplex to an acceptor RNA template (right) compared to that of GsI-IIC RT bound to a continuous RNA template/DNA primer heteroduplex (left; PDB: 6AR1). Protein regions: fingers (salmon), insertions (red), palm (dark blue), thumb (green), D domain (gold), acceptor RNA template (yellow), donor RNA template (purple), DNA primer (cyan), dATP (black). T and P denote the template and primer strands respectively. The RT0 loop is highlighted in a dotted circle. *B*, schematic of protein-nucleic acid interactions in the template-switching structure. Nucleotide positions are denoted as template (T) or primer (P) strand numbered from the templating nucleotide at the RT active site (T-1). Interactions between nucleic acid and amino acid residues are indicated by a black line (polar interaction), dotted line (non-polar interaction), or a double-headed arrow (RNA 2′ OH H-bond). Other color codes are as in panel *A*. The first nucleotide of the acceptor (the U at T-4) could not be modeled and is shown in a lighter shade of yellow. *C*, amino acid sequence alignments of the RT0 loop (top) and fingertips loop (bottom) regions of group II intron RTs (red; GsI-IIC RT (E2GM63), *Roseburia intestinalis* (D4L313), *Eubacterium rectale* (D4JMT6), Ll.LtrB (P0A3U0), TeI4h (Q8DMK2), *Saccharomyces cerevisiae* aI2 (P03876)) and non-LTR-retroelement RTs (black; human LINE-1 (Ll) orf2p (O00370), R2 *Bombyx mori* (V9H052), Jockey *Drosophila funebris* RT (P21329)). Uniprot IDs are indicated in parentheses. Protein sequences were aligned using the MAFFT algorithm and colored using Clustalx settings.

The complex crystallized in the space group C2, with the asymmetric unit containing two pseudo-symmetric RT monomers, each bound to the acceptor RNA and donor duplex. We determined initial phases by molecular replacement, using the previously determined GsI-IIC RT structure as the search model (PDB: 6AR3; (24)) with the dATP and parts of the template strand removed to reduce model bias. Similar to the previously solved GsI-IIC RT structure, we encountered problems of crystal twinning, discussed in detail in the prior work (24). We were able to refine the structure to *R_Work_* and *R_Free_* values of 27.1% and 32.4% at 3.2-Å resolution with other collection and refinement statistics summarized in Table 1. We were unable to crystallize the complex in the absence of dATP or with donor duplexes having a blunt-end or mismatched 3’-overhang nucleotide. These findings suggest that both base pairing of the 3’ nucleotide of the acceptor to the 1-nt 3’-overhang nucleotide of the DNA oligonucleotide and binding of a dNTP complementary to the templating RNA base are required for the RT to form a crystallizable complex.

**Table 1.**
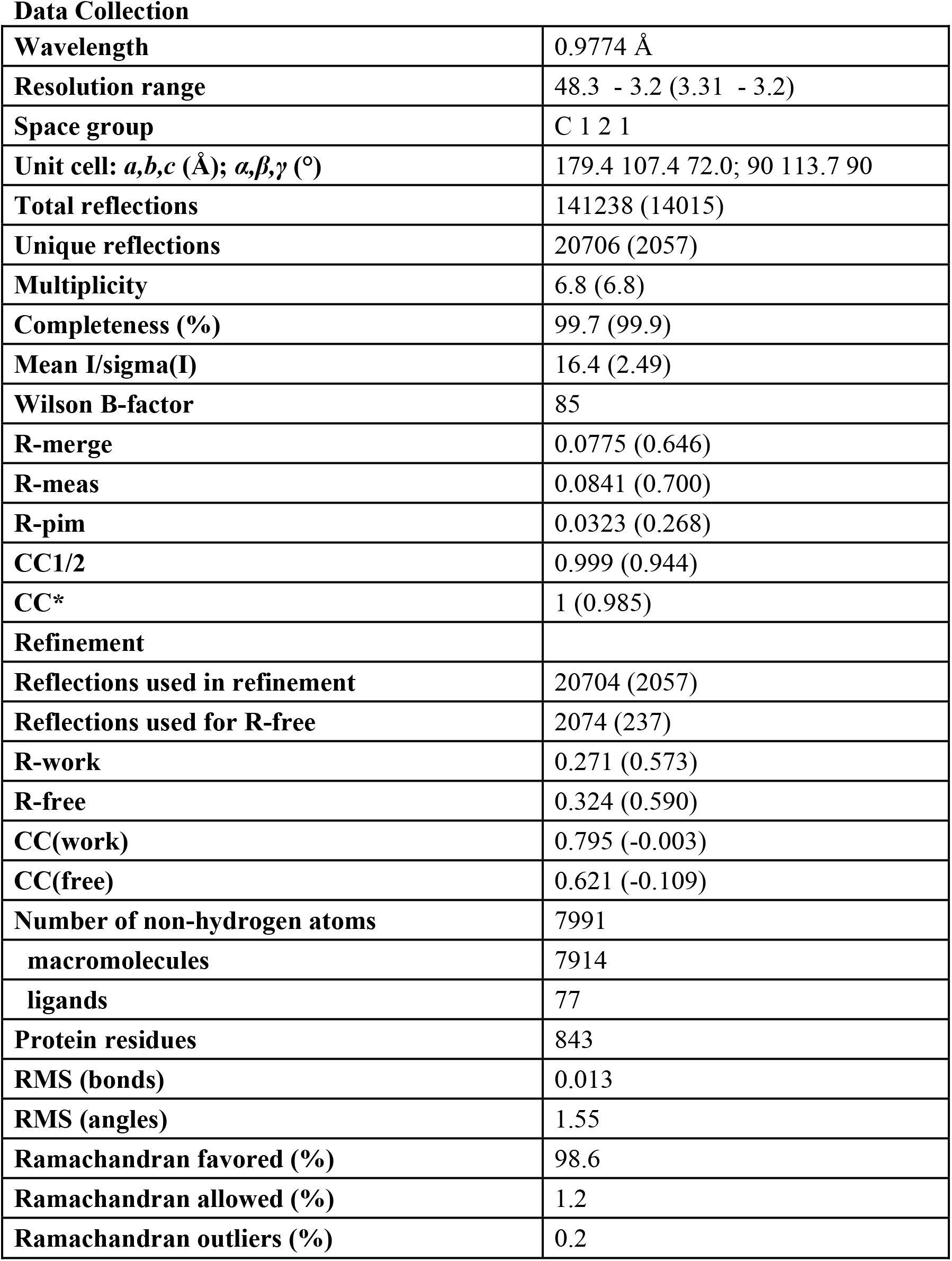
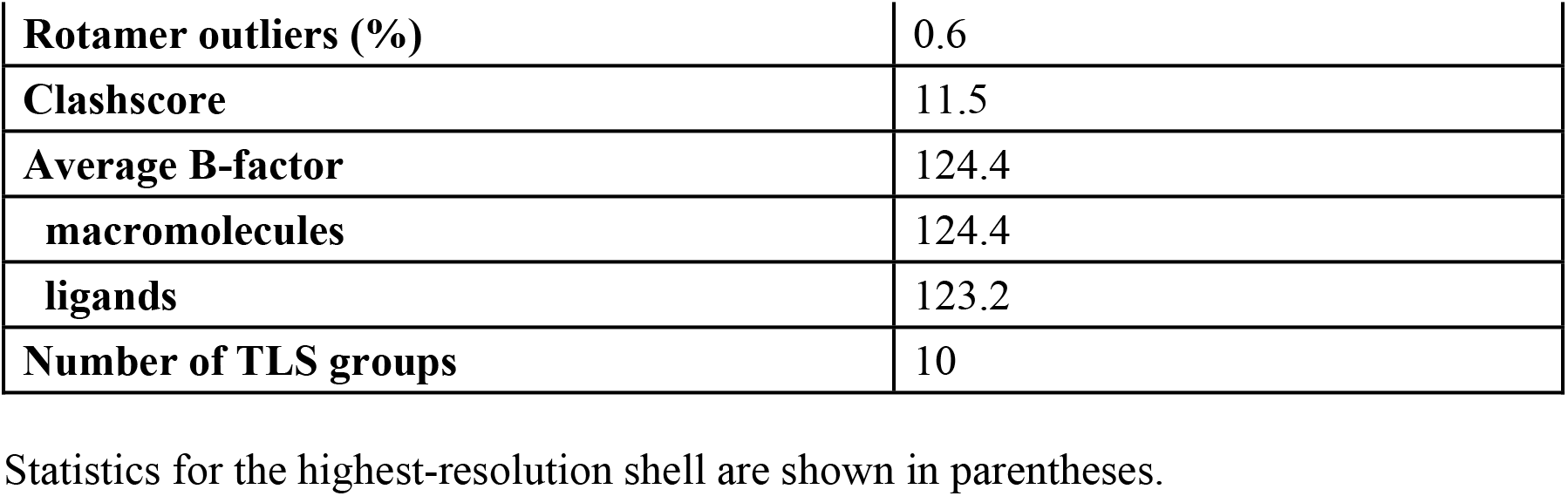
Collection and refinement statistics for the GsI-IIC RT template-switching complex structure.

The overall conformation of GsI-IIC RT in the template-switching complex was largely the same as that bound to a continuous RNA template (PDB: 6AR1 (24)) (Fig. 1*A*). The protein follows the canonical hand-like fold of other RTs with fingers, palm and thumb regions, plus an appended DNA-binding domain (D) that contributes to DNA target site recognition during group II intron mobility ("retrohoming") to a new DNA site (Fig. 1*A*) (2). The last three nucleotides of the acceptor RNA (T+1 to T-2) sit within a pocket that is formed by the NTE and the fingertips loop and would ordinarily be occupied by upstream regions of a continuous RNA template (Fig. 2). T-3, which lies outside the template-switching pocket, is ordered as a result of a crystal contact to a symmetry-related GsI-IIC RT thumb domain, while the 5’ nucleotide of the acceptor (T-4), which lies farther outside the template-switching pocket, could not be modeled and is likely disordered.

**Figure 2.**
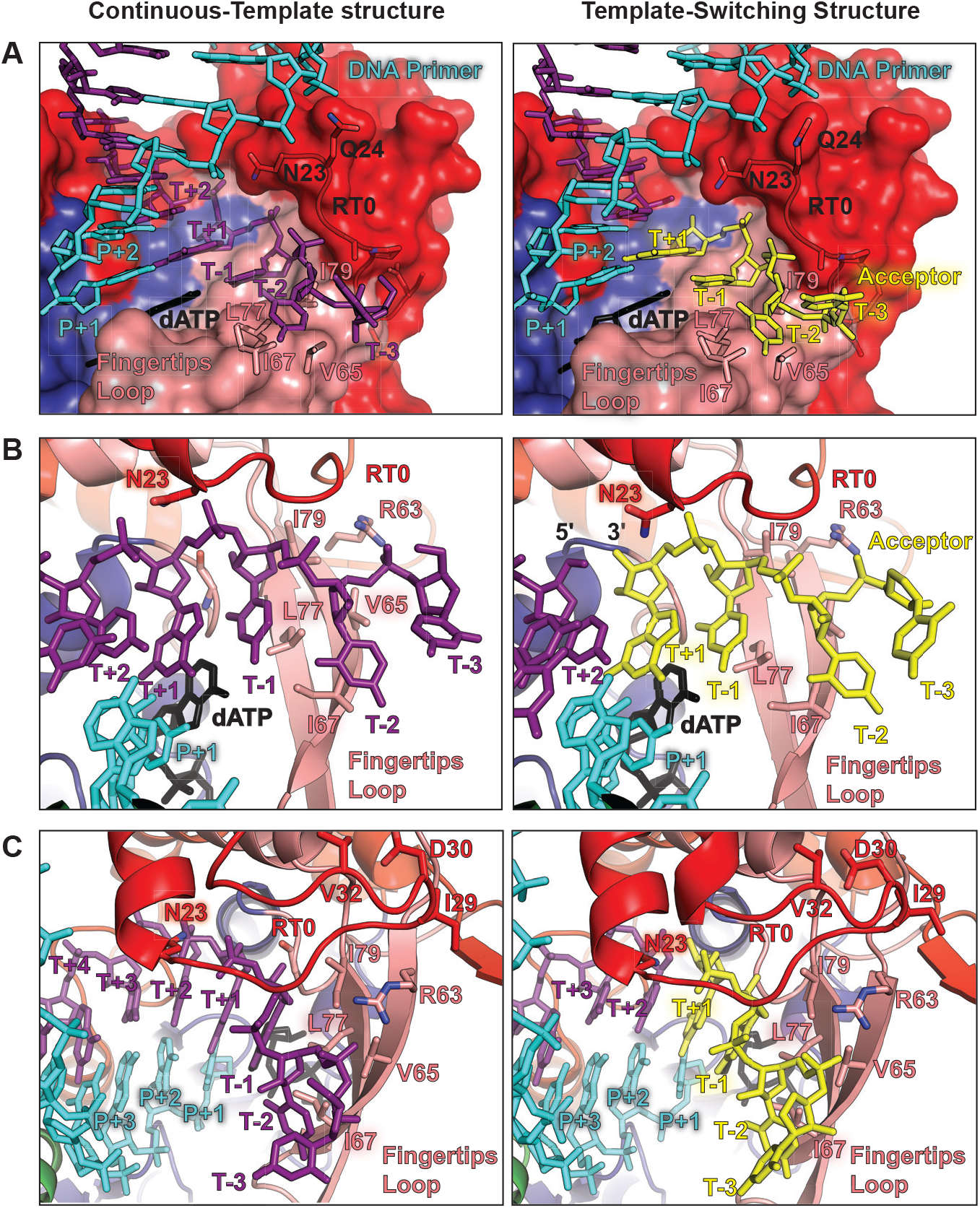
Binding of the 3’ end of the acceptor RNA within the template-switching pocket. *A*, close-up views comparing the binding of the 3’ end of the acceptor within the template-switching pocket formed by the NTE and fingertips loop (right) with the binding of a continuous RNA template in the same region of the protein (left; PDB: 6AR1). Protein regions: fingers (salmon), insertions (red), palm (dark blue); acceptor RNA template (yellow), donor RNA or continuous template (purple), DNA primer (cyan), dATP (black). Nucleic acids are depicted as sticks, and the protein is depicted in surface-filling representation with some residues highlighted as sticks. *B*, comparison of the same structures with rotation to give a better view of the junction region between the donor and acceptor RNA templates. Nucleic acids and protein regions are colored as in panel *A*. Nucleic acids are in stick representation and protein is in cartoon representation with some residues highlighted as sticks. *C*, comparisons of the same structures with rotation to show the binding of the 3’ end of the acceptor beneath the ‘lid’ of the RT0 loop within the template-switching pocket. Nucleic acids and protein regions are colored and shown in the same representation as in panel *B*.

In agreement with the template-switching mechanism inferred from biochemical analysis (27), the 3’ G of the acceptor RNA at T+1 is base-paired to the 3’-overhang nucleotide of the DNA primer (P+1) at the position corresponding to a newly formed base pair after translocation out of the RT active site, while the penultimate nucleotide of the acceptor at T-1 occupies the position of the templating base and is base paired to the incoming dATP at the RT active site (Fig. 1*A*). The high specificity of template switching likely reflects that these two base-pairing interactions occur sequentially, with the second between the templating base and incoming dNTP accompanied by a conformational change required for the initiation of reverse transcription, which drives the reaction forward (see Discussion).

Most of the contacts between GsI-IIC RT and the acceptor RNA and donor duplex are indistinguishable at this resolution from those for the RNA template and annealed DNA primer in the continuous template structure (Fig. 1*B*) (24). The 3’ end of the RNA acceptor at T+1, which is base-paired to the 1-nt DNA overhang, and the templating base at T-1 rest near the fingers/palm junction on an inner surface of the protein formed by residues F110, G192 in the PQG loop, G193, and P194, with hydrogen bonds from the backbone carbonyls of G192 and G193 to the 2’-OHs of T-1 and T+1, respectively (Fig. 1*B*).

Near the junction between the donor and acceptor RNAs, the RT0 loop of the NTE is in the same conformation as in the continuous RNA template structure (Fig. 2*A*; RMSD = 0.851 Å). The N-terminal portion of the loop from residues N23 to A26 (Fig. 1*C*) forms a lid over the phosphate backbone of T+1 and T-1, with hydrogen bonds from the side chain of N23 to the T+1 phosphate and from the backbone amide of A26 to the T-1 phosphate (Fig. 2*A*). As in other structures of group II intron RTs, including high-resolution crystal structures of N-terminal group II intron RT fragments without bound nucleic acids, the RT0 loop is well defined and anchored to the body of the protein by conserved hydrophobic residues at the central tip and C-terminal end of the loop (I29 and V32, respectively) and by hydrogen bonds between the side chain of R85 in the fingers and the backbone carbonyls of G25 and A26 in the loop (24,28–30) (Fig. 1*B* and Fig. 2*C*). A highly conserved aspartate residue (D30) hydrogen bonds to the nearby G31 backbone amide (Fig. 2*C*). It may also form a hydrogen bond to T81 of the fingers, but this interaction is not conserved in other group II intron RT structures (28–30).

In the continuous template structure, N23 at the beginning of the RT0 loop, rests midway between the two phosphate groups on either side of T+1, which corresponds to the 3’ nucleotide of the acceptor. In the template-switching structure, however, where T+1 is separated from the 5’ nucleotide of the donor (T+2) by a gap lacking a phosphate group, N23 shifts away from this position to suggest a closer interaction with the phosphate between T-1 and T+1, to which R85 from the fingers hydrogen bonds on the opposite side (Fig. 1*B*, and Figs. 2, *A* and *B*). The adjacent residue, Q24, interacts with the phosphate of the primer strand between positions P+6 and P+7, potentially helping to anchor the donor duplex near the template-switching junction.

The remaining binding surface for the acceptor RNA within the template-switching pocket is provided by the fingertips loop, a conserved structural feature found in all RTs (Fig. 2*, B* and *C*). The fingertips loop consists of two anti-parallel *β* strands, which are hinged and upon dNTP binding move the side chains of positively charged residues (R75 and K69 in GsI-IIC RT) into the RT active site to properly position the dNTP substrate and stabilize the negative charges of the triphosphate (31, 32). In the template-switching structure, the fingertips loop is closed over the RT active site in the same conformation as in the continuous template structure (RMSD of 0.684 Å; Fig. 2*C*). The opposite face of the fingertips loop contains a series of conserved branched hydrophobic residues that cradle the underside of the RNA acceptor at T-1 and T-2, with L77 and I79 contacting the T-1 base and ribose, respectively, and V65 and I67 contacting the T-2 ribose and base, respectively (Fig. 2, *B* and *C*; see also Fig. 5*A* below). R63, which is located at the base of the fingertips loop, forms a polar contact to the phosphate connecting T-2 and T-3, representing the 5’-most interaction of the RT with the acceptor RNA in the template-switching pocket, and P68, which sits near the end of the fingertips loop between the two *β* strands, makes a distant non-polar interaction with the base of T-2 (Fig. 1*B* and Fig. 2, *B* and *C*; see also Fig. 5*A* below). The contacts between the six fingertips loop residues identified as binding the acceptor RNA were not detectably changed when compared to the continuous template structure (24). These six residues together with N23 of the NTE are often conserved in group II intron RTs (Fig. 1*C*) and potentially represent important contacts for template switching (22).

### Template switching of wild-type GsI-IIC RT and effect of mutations in the RT0 loop

To investigate the functions of the amino acid residues identified in the crystal structure as making side-chain contacts with the acceptor RNA, we constructed a series of GsI-IIC RT mutants and compared their template-switching and primer extension activities to those of the wild-type (WT) enzyme. Since we were interested in mutations that might affect binding of the acceptor, we first examined the template-switching activity of the WT enzyme as a function of acceptor RNA concentration (Fig. 3). To monitor binding of the acceptor and subsequent nucleotide incorporation, we used saturating dNTP concentrations with varying concentrations of a 21-nt acceptor RNA with a 3’-C residue and a fixed concentration of a donor RNA/DNA heteroduplex with a complementary 1-nt 3’-G DNA overhang (20 nM, substoichiometric relative to enzyme). Except for the fixed 1-nt 3’-G overhang, the donor duplex used for biochemical experiments corresponds to a version used for TGIRT-seq and is unrelated to the shorter donor duplex used for crystallization (see Fig. 3 legend and Table S1).

**Figure 3.**
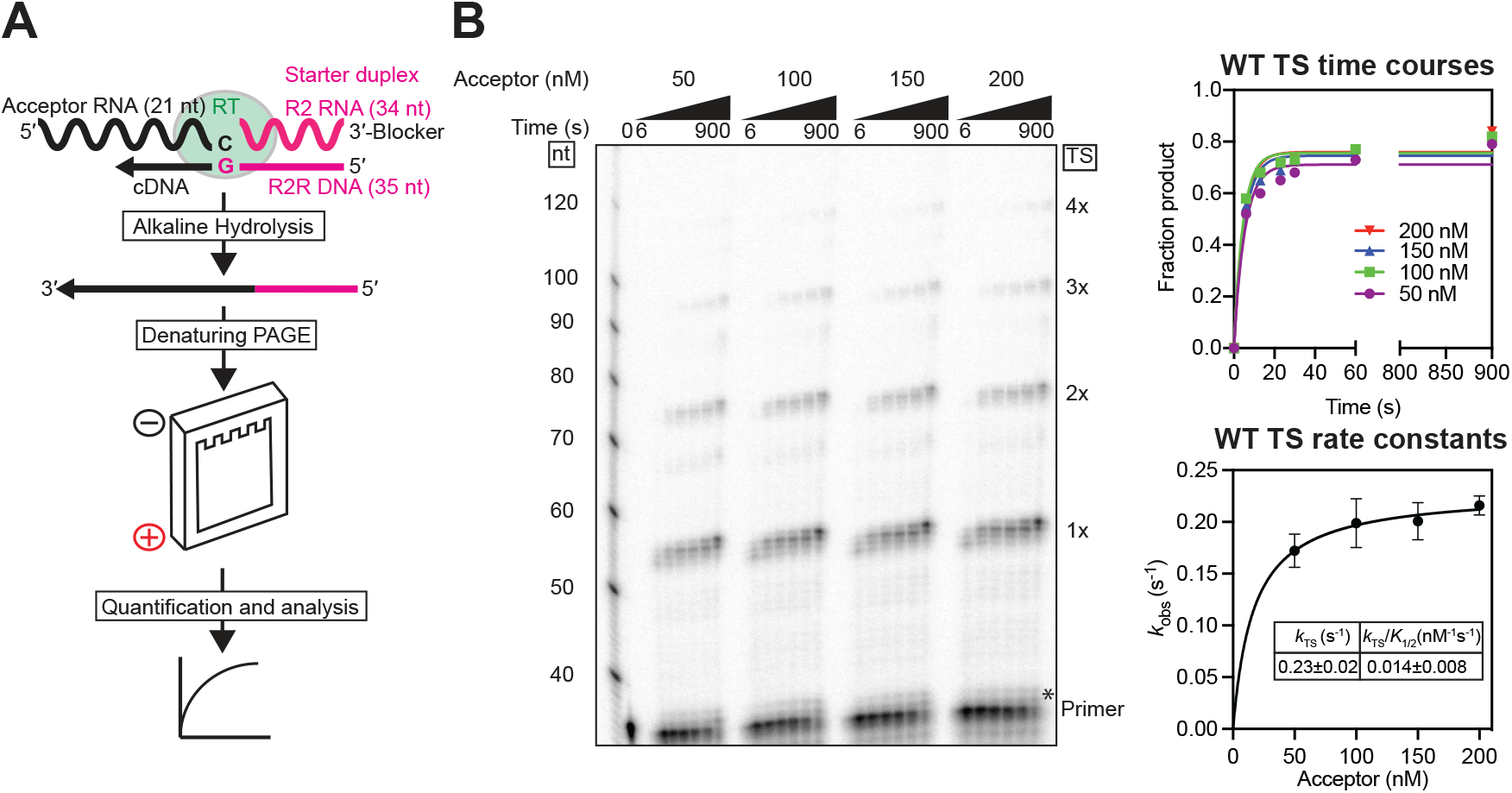
Overview of template-switching assays and determination of saturating acceptor RNA concentrations. *A*, outline of template-switching assay. GsI-IIC RT (green oval) was pre-incubated with a starter duplex (magenta) consisting of a 34-nt RNA oligonucleotide containing an Illumina Read 2 (R2) sequence annealed to a complementary 5′-^32^P-labeled 35-nt DNA primer (R2R) leaving a 1-nt 3′-DNA overhang (nucleic acid sequences in Table S1). The 3′-DNA overhang nucleotide (G) base pairs with the 3′ nucleotide (C) of a 21-nt acceptor RNA (black) for template switching, leading to the synthesis of a full-length cDNA of the acceptor RNA with the R2R oligonucleotide linked to its 5′ end. After incubation with NaOH to degrade RNA and neutralization with equimolar HCl, the cDNAs resulting from template switching were analyzed by electrophoresis in a denaturing 6% polyacrylamide gel, which was dried and quantified with a phosphorimager. *B*, determination of saturating acceptor RNA concentrations. Time courses of template-switching (TS) reactions using 200 nM wild-type (WT) GsI-IIC RT, 20 nM donor RNA template/DNA primer duplex (5′-^32^P-labeled on DNA primer), indicated concentrations of a 21-nt acceptor RNA template, and 4 mM dNTPs (an equimolar mix of 1 mM dATP, dCTP, dGTP, and dTTP) in reaction medium containing 200 mM NaCl at 60 °C. Aliquots were quenched at times ranging from 6 to 900 s, and the products were analyzed by denaturing PAGE. The Figure shows a representative gel from one of three repeats of the experiment. The numbers to the left of the gel indicate size markers (a 5′-^32^P-labeled single-stranded DNA ladder; ss20 DNA Ladder, Simplex Sciences) run in a parallel lane, and the labels to the right indicate products resulting from the initial template switch (1x) and subsequent end–to–end template switches from the 5′ end of one acceptor to the 3′ end of another (2x, 3x, etc.). The asterisk at the bottom right of the gel indicates the position of bands resulting from NTA to the 3′ end of the DNA primer. The plot at the upper right shows time courses of production of template-switching products (*i.e.*, products >2 nt larger than the primer) at each RNA acceptor concentration. The plot at the bottom right shows *k*_obs_ as a function of acceptor concentration fit by a hyperbolic equation to obtain the maximal rate constant *k*_TS_ and the second-order rate constant *k*_TS_/*K*_1/2_, with the error bars in the plot showing the standard error of the mean for three repeats of each time course and the uncertainties in the inset table indicating the standard error of the fit.

The template-switching products in these experiments showed a characteristic ladder of bands that differ by 21 nt, reflecting multiple sequential template switches from the 5’ end of one RNA template to the 3’ end of another (Fig. 3*B*). Reaction time courses revealed a single predominant kinetic phase for the appearance of template-switching products, giving an observed rate constant (*k*_obs_) for each acceptor RNA concentration (Fig. 3*B*). A fit of the observed rate constant as a function of acceptor concentration gave a maximal rate constant for template-switching (*k*_TS_) of 0.23 ± 0.02 s^-1^, in agreement with previous measurements done at saturating acceptor concentrations (0.25 s^-1^ (27)). The observed rate constant approached this maximum value even with the lowest concentration of acceptor (50 nM) that could be tested while maintaining the acceptor in excess of the GsI-IIC RT-starter duplex complex. Within this limited concentration-dependent regime, the fit gave an estimated second-order rate constant (*k*_TS_/*K*_1/2_) of 0.014 nM^-1^ s^-1^ ± 0.008. The latter value, also known as the specificity constant, represents the rate constant for acceptor binding multiplied by the probability of subsequent chain extension and was within 2 to 3 orders of magnitude of the diffusion limit, underscoring the efficiency of GsI-IIC RT for template switching (Fig. 3*B*).

We previously compared the template-switching activities of WT and several mutant GsI-IIC RTs with alterations in conserved residues by using the same combination of a 3’-C acceptor RNA and 1-nt 3’-G DNA overhang donor duplex but in reactions incubated for a fixed time (15 min) at saturating concentrations of acceptor RNA and dNTPs (24). We found that mutations in a conserved anchoring residue in the RT0 loop (I29R, Fig. 1*C*) and in a conserved arginine in the fingers whose side chain H-bonds to a backbone carbonyl in the RT0 loop and the phosphate between T-1 and T+1 of the acceptor (R85A, Fig. 1*C*) strongly inhibited template switching while retaining high primer extension activity (24). By contrast, the mutation D30A, in another conserved residue that anchors the RT0 loop to the body of the protein, had no detectable effect on either template switching or primer extension (24). Additionally, the RT0 loop mutant 23-28/6G, in which the six residues at positions 23-28 were replaced by six glycines, retained high template-switching activity, while the mutant 23-31/4G, with a shorter RT0 loop in which residues 23-31 were replaced by four glycines, substantially inhibited template-switching activity (24). These findings suggested that the size and/or conformation of the RT0 loop might significantly impact template switching (24).

Based on the above results and the crystal structure of the template-switching complex, we focused more detailed analysis on N23 and Q24, two solvent exposed RT0 loop residues with prominent polar side-chain interactions with the acceptor backbone and donor duplex, respectively, and the RT0 loop mutant 23-31/4G, which inhibited template-switching activity in the previous assays (24). N23 was of interest as a conserved residue (Fig. 1*C*) whose side chain sits between the phosphates on either side of T+1 in the continuous-template structure but shifts closer to the upstream phosphate in the template-switching structure, where a downstream phosphate is not present in the gap between the donor and acceptor (Fig. 2*B*). This finding raised the possibilities that the N23 side chain could toggle between the two phosphates and that its interaction with the upstream phosphate could be particularly important for binding an acceptor RNA during template switching, where there is a discontinuity in the phosphate backbone. The adjacent residue Q24 makes a side-chain polar contact with the phosphate between P+6 and P+7 on the primer strand, potentially anchoring the donor duplex near the template-switching junction (see above).

Nevertheless, we found that the mutants N23A, Q24G, N23A/Q24A and N23G/Q24G had no effect on primer extension activity (Fig. S1) and displayed near-saturating kinetics at all the acceptor RNA concentrations tested, giving maximal rate constants of 0.28-0.32 s^-1^ and *k*_TS_/*K*_1/2_ values of ∼0.01 to 0.02 nM^-1^ s^-1^, indistinguishable from those of WT (Table 2, Fig. 4, and Fig. S2). These results indicate that the side chains of N23 and Q24 do not play major roles in acceptor RNA binding or chain extension. It remains possible that Q24 stabilizes binding of the duplex, as the experiments were performed using a concentration of enzyme sufficient to saturate duplex binding.

**Figure 4.**
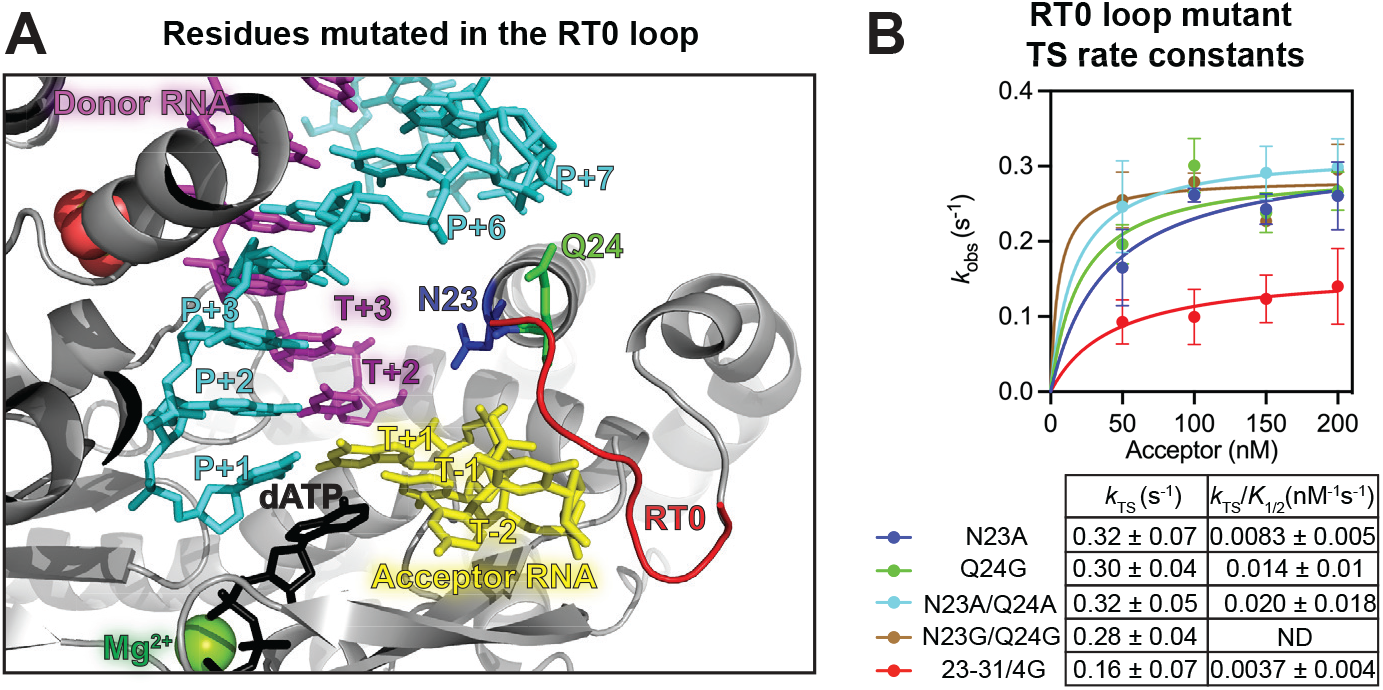
The effect of RT0 loop mutations on template switching by GsI-IIC RT. *A*, closeup view of the RT0 loop highlighting the location of mutated amino acid residues N23 (blue stick) and Q24 (green stick) and the polypeptide backbone of residues 23-31 (red cartoon). Nucleic acids are colored as in Fig. 1A. *B*, the plot shows *k*_obs_ as a function of acceptor concentration fit by a hyperbolic equation to obtain the maximal rate constant *k*_TS_ and the second-order rate constant *k*_TS_/*K*_1/2_. Template-switching reactions were done as described in Fig. 3. Each rate constant measurement was performed twice and representative time courses are shown in Fig. S2. The error bars in the plot show the standard error of the mean, and the uncertainties in the *k*_TS_ and *k*_TS_/*K*_1/2_ values in the table below indicate the standard error of the fit.

**Table 2.** Summary of kinetic parameters determined for wild-type and mutant GsI-IIC RTs. k_TS_ and *k*_TS_ /*K*_1/2_ values are the best fit ± standard errors of the fit values obtained from fitting of a hyperbolic equation to the observed rate constants from two to three repeats of the time courses in the experiments of Figs. 3, 4, 5, and 6 (see Experimental procedures). *K*_1/2_ values, which were not precisely determined by the time course data due to saturation of the observed rate constants at low concentrations of acceptor for the WT and most of the mutants, are approximate values obtained from these fits, with relative uncertainties comparable to those for *k*_TS_/*K*_1/2._ PE nt/s values are approximate mean values obtained from the mid-point of the 10 s time point in two to three repeats of the time courses in Fig. S1 Abbreviations: NA, not applicable; ND, not determined; PE, primer extension.

| Enzyme | Mutated region | $k_{TS}$ (s<sup>-1</sup>) | $k_{TS} / K_{1/2}$ (nM<sup>-1</sup> s<sup>-1</sup>) | $K_{1/2}$ (nM) | PE (nt/s) |
| --- | --- | --- | --- | --- | --- |
| **WT** | NA | 0.23 ± 0.02 | 0.014 ± 0.008 | ~ 17 | ~20 |
| **N23A** | RT0 loop | 0.32 ± 0.07 | 0.0083 ± 0.005 | ~ 39 | ~20 |
| **Q24G** | RT0 loop | 0.30 ± 0.04 | 0.014 ± 0.01 | ~ 21 | ~20 |
| **N23A/Q24A** | RT0 loop | 0.32 ± 0.05 | 0.020 ± 0.018 | ~ 16 | ~20 |
| **N23G/Q24G** | RT0 loop | 0.28 ± 0.04 | ND | ND | ~20 |
| **23-31/4G** | RT0 loop | 0.16 ± 0.07 | 0.0037 ± 0.004 | ~ 44 | ~20 |
| **R63A** | Fingertips loop | 0.21 ± 0.06 | 0.0084 ± 0.01 | ~ 25 | ~20 |
| **V65A/I67A** | Fingertips loop | 0.25 ± 0.05 | 0.0077 ± 0.005 | ~ 33 | ~15 |
| **L77A/I79A** | Fingertips loop | ND | 0.00040 ± 0.0002 | ND | ~15 |
| **F143A** | dNTP-binding site | 0.19 ± 0.03 | 0.0034 ± 0.001 | ~ 56 | ~10 |

The movement of the side chain of the N23 away from the gap between the donor and acceptor in the template-switching structure also led us to wonder whether adding a 5′ phosphate to the RNA donor would impact template switching. To test this idea, we performed template-switching reactions with WT and N23A with the same acceptor RNA and either 5′ OH and 5′ phosphate donors (Fig. S3). For both proteins, template-switching activity with the 5′ phosphate donor decreased, likely due to steric clashes of the additional oxygen on the 5′ phosphate of donor compared to a phosphodiester bond (Fig. S3).

Finally, the mutation 23-31/4G had no detectable effect on primer extension activity (Fig. S1) but strongly inhibited template-switching activity, with the mutant showing subsaturating kinetics over the range of acceptor RNA concentrations tested with a decreased *k*_TS_ = 0.16 s^-1^ and a 3.5-fold lower *k*_TS_/*K*_1/2_ than the wild-type enzyme (Fig. 4, Fig. S2, and Table 2) (24). These results indicate that the altered RT0 loop conformation, resulting from shortening of the loop and/or loss of the I29 anchoring interaction (see above), substantially inhibited binding of the acceptor RNA. The lack of effect of this mutation on primer extension indicates that when the template-switching pocket is occupied by a continuous RNA template the shorter, glycine-rich RT0 loop is capable of adopting a conformation that does not impede reverse transcription.

### Template switching of GsI-IIC RTs with mutations in the fingertips loop

The remaining residues that make potentially critical side-chain contacts with the acceptor RNA are located in the fingertips loop, a conserved structural feature that forms part of the dNTP-binding pocket and contacts parts of the DNA primer and RNA acceptor. Here, we focused on five of the residues identified in the crystal structure as making side-chain contacts with acceptor RNA: R63, which is located at the base of the loop and makes a side-chain contact with the phosphate between T-2 and T-3; V65 and I67, which are located centrally in the first β-strand of the hairpin with their hydrophobic side chains engaged in non-polar interactions with the ribose and base of T-2, respectively; and L77 and I79, which are located on the anti-parallel *β*-strand with their side chains making non-polar contacts with the sugar and base of T-1, the templating nucleotide at the RNA active site, and with the side chain of I79 also interacting with T-2 (Fig. 1 and Fig. 5). The mutation P68A is the sixth fingertips loop residue that makes a side-chain contact with the acceptor (a distant non-polar contact paper with the base of T-2) did not inhibit template switching in initial assays (Fig. S4*B*) and was not subjected to more detailed analysis.

**Figure 5.**
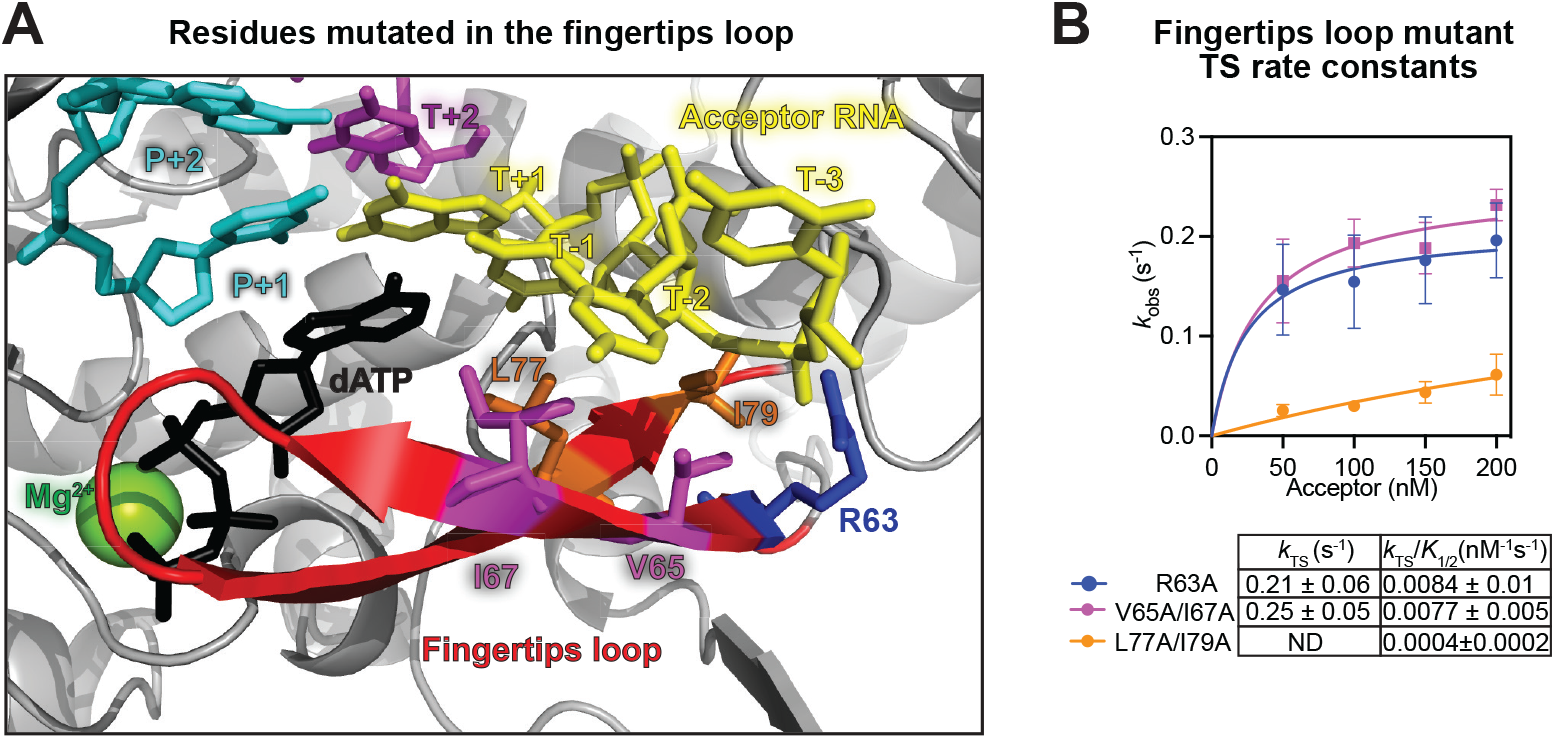
The effect of fingertips loop mutations on template switching by GsI-IIC RT. *A*, closeup view of the fingertips loop (red cartoon) highlighting the location of mutated amino acid residues R63 (blue stick), V65/I67 (magenta stick), and L77/I79 (orange stick). Nucleic acids are colored as in Fig. 1A. *B*, the plot shows *k*_obs_ as a function of acceptor concentration fit by a hyperbolic equation to obtain the maximal rate constant *k*_TS_ and the second-order rate constant *k*_TS_/*K*_1/2_. Template-switching reactions were done as described in Fig. 3, with each rate constant measurement performed twice and representative time courses shown in Fig. S4. The error bars in the plot show the standard error of the mean, and the uncertainties in the *k*_TS_ and *k*_TS_/*K*_1/2_ values in the table below indicate the standard error of the fit.

To assess the contributions of the other fingertips loop residues to template switching, we constructed the mutant R63A and double mutants V65A/I67A and L77A/I79A. The double mutant L77A/I79A had the strongest effect on template switching, with a *k*_TS_/*K*_1/2_ value (0.0004 ± 0.0002 nM^-1^ s^-1^; Fig. 5*B* and Fig. S4) that was 25-fold lower than that for WT (Fig. 4*B* and Table 2). R63A and V65A/I67A have *k*_TS_ values similar to wild type, but lower *k*_TS_/*K*_1/2_ values (0.0084 ± 0.01 and 0.0077 nM^-1^s^-1^ ± 0.005 compared to wild type at 0.014 nM^-1^s^-1^; Table 2 and Figs. 3 and 5), most likely reflecting impaired binding of the acceptor RNA. All of these mutants retained high primer extension activity (15 to 20 nt/sec; Table 2 and Fig. S1), suggesting that the fingertips loop interactions are more critical for template switching than for reverse transcription on a continuous RNA template.

### Identification of an amino acid residue that is critical for non-templated nucleotide addition

Our previous kinetic analysis indicated that template switching occurs largely by an ordered process in which a single non-templated nucleotide is added to the 3′-end of a completed cDNA, thereby creating a 1-nt 3′-DNA overhang that can then base pair to the 3′ end of an acceptor RNA (27). To further investigate the contribution of NTA to template switching, we focused on F143, which is located in the dNTP binding pocket and may contribute to the selectivity of GsI-IIC RT for dNTPs by sterically blocking the 2’ OH present in rNTPs (24, 33). As in other RTs, the aromatic side chain of F143 is aligned for a pi stacking interaction with the deoxyribose sugar of the incoming dNTP (Fig. 6*A*), and we hypothesized that this stabilizing interaction could be important for NTA, during which the added dNTP is not base paired with a templating nucleotide.

**Figure 6.**
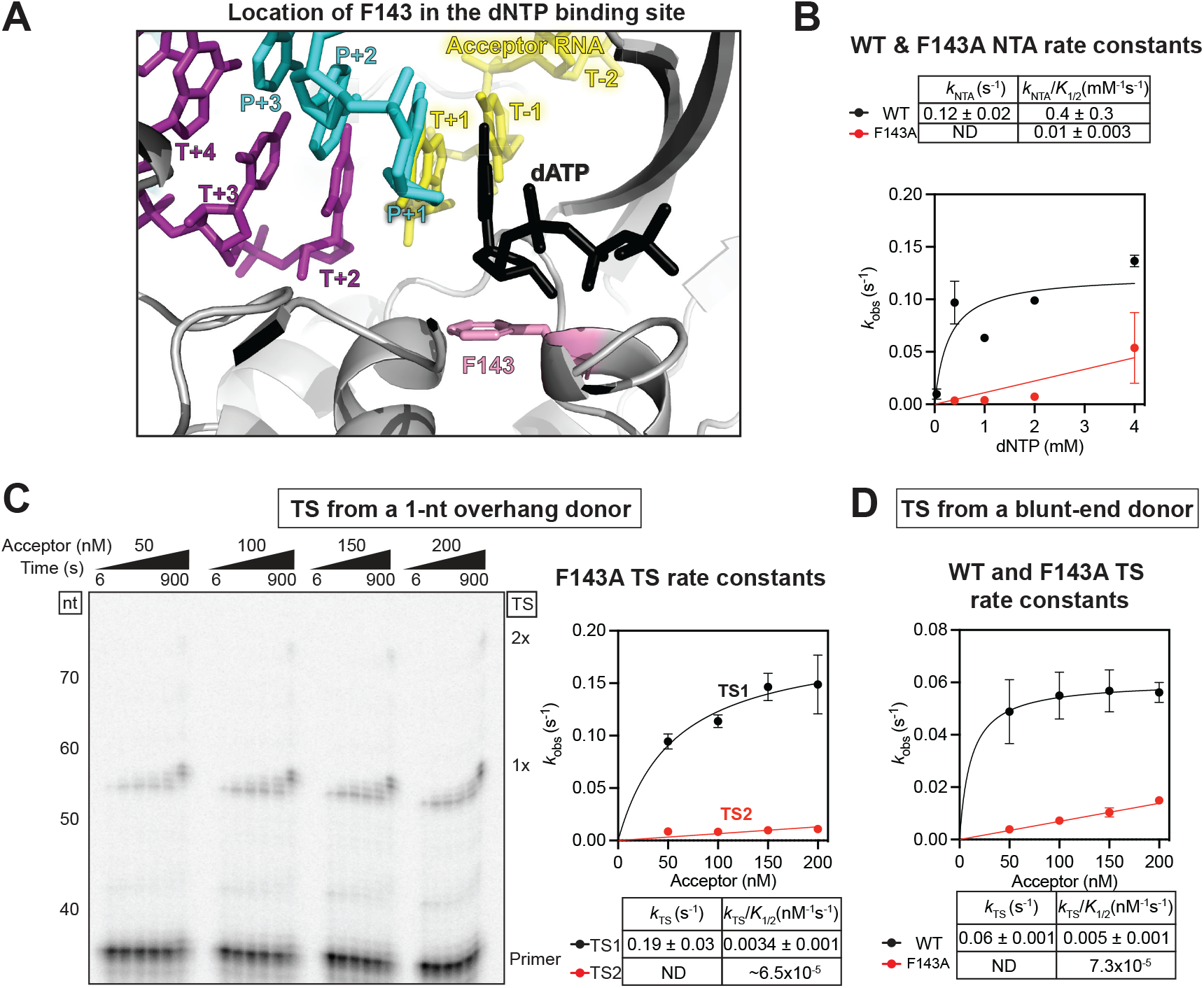
The effect of the F143A mutation on non-templated nucleotide addition and template switching by GsI-IIC RT. *A*, closeup view of the dNTP-binding site showing F143 (pink stick) pi stacking with the ribose of the incoming dATP (black stick). Nucleic acids are colored as in Fig. 1A. *B*, NTA reactions for WT and F143A mutant GsI-IIC RT as a function of dNTP concentrations. The reactions were performed with blunt-end RNA template/DNA primer duplex of the same sequence as that used for template switching but lacking the 1-nt 3’-DNA overhang and under the same reaction conditions as those used for template switching in Fig. 3. The plot shows *k*_obs_ as a function of dNTP concentration fit by a hyperbolic equation to obtain the maximal rate constant *k*_NTA_ and the second-order rate constant *k*_NTA_/*K*_1/2_ values. Each rate constant determination was performed at least twice, and a representative gel is shown in Fig. S5*A*. The error bars in the plot show the standard error of the mean, and the uncertainties in the *k*_NTA_ and *k*_NTA_/*K*_1/2_ values in the table above indicate the standard error of the fit. The 0.04 mM concentration for F143A was omitted because of poor fitting due to low amounts of product formation. *C*, template-switching reactions of the F143A mutant from a 1-nt 3’-DNA overhang RNA template/DNA primer duplex. Reactions were performed as described in Fig. 3. The numbers to the left of the gel indicate the length and position of size markers (5′-^32^P-labeled single-stranded DNA ladder; ss20 DNA Ladder, Simplex Sciences) run in a parallel lane of the same gel. Each rate constant determination was performed at least twice. The plot to the right shows the observed rate constant *k*_obs_ as a function of acceptor RNA concentration for the initial template switch from the donor duplex (TS1) and the second template switch from the 5’ end of first RNA template to the 3’ end of a second RNA template (TS2). The data points were fit by a hyperbolic function. *D*, template switching of WT and F143A mutant GsI-IIC RT from a blunt-end duplex. Template-switching reactions were done as described in Fig. 3. Each rate constant determination was performed three times, and a representative gel is shown in Fig. 5B. The plot shows *k*_obs_ as a function of acceptor concentration fit by a hyperbolic equation to obtain the maximal rate constant *k*_TS_ and the second-order rate constant *k*_TS_/*K*_1/2_, as indicated in the inset table. In panels *C* and *D*, the error bars in the plots show the standard error of the mean, and the uncertainties in the tables below each plot indicate the standard error of the fit.

We constructed the mutant F143A and measured its kinetics of NTA to a blunt-end starter duplex in the absence of an acceptor RNA. We found that NTA was strongly diminished, with a ∼ 40-fold lower *k*_NTA_/*K*_1/2_ value than the WT enzyme (Fig. 6*B*, Fig. S5*A*). The decrease in rate was smaller at the highest dNTP concentrations measured, indicating that the decreased *k*_NTA_/*K*_1/2_ value is primarily due to weaker binding of the incoming dNTP (Fig. 6*B* and Fig. S5*A*). The F143A mutant retained substantial primer extension activity, synthesizing full-length cDNAs of a 1.1-kb RNA template with no indication of premature stops and a relatively small decrease in rate (10 nt/s compared to 20 nt/s for WT at 4 mM dNTPs; Fig. S1 and Table 2).

Next, we measured the template-switching activity of F143A with a donor RNA template/DNA primer heteroduplex with a pre-formed 1-nt 3’-DNA overhang and increasing concentrations of acceptor RNA under our standard reaction conditions with 4 mM dNTPs. We found that the F143A mutant produced cDNA products at a rapid rate (*k*_TS_ = 0.19 ± 0.03 s^-1^), reflecting that the first template switch from the donor duplex (TS1) was relatively unaffected, but with greatly reduced efficiency for the second template switch (TS2; *k*_TS_/*K*_1/2_ = ∼ 6.5 x 10^-5^; compare second template switch by F143A in the gel of Fig. 6*C* to WT in Fig. 3*B* and see below).

Finally, we anticipated that template switching from a blunt-end donor would be strongly decreased by the F143A mutation with kinetic parameters similar to that of its second template switch from a 1-nt overhang substrate. As seen in Fig. 6*D*, this was indeed the case, with template switching from a blunt-end duplex by F143A showing an ∼70-fold decrease in *k*_TS_/*K*_1/2_ compared to WT (*k*_TS_/*K*_1/2_ of 0.005 ± 0.001 for WT and 7.3 x 10^-5^ nM^-1^ s^-1^ for F143A; Fig. S5*B*, Table 2). Collectively, our results show that the F143A mutation had at most a small effect on the first template switch from a donor RNA template/DNA primer heteroduplex with 4 mM dNTPs, but strongly inhibited multiple end-to-end template-switches, which depend upon NTA to produce a single-nucleotide overhang (27).

Together, our results indicate that the F143A mutation weakens binding of an unpaired dNTP at the RT active site, thereby decreasing NTA activity. Although the F143A mutation may also weaken dNTP binding during cDNA synthesis, there was no decrease in the amplitude and only a relatively small decrease (two-fold) in the rate (nt/s) of primer extension at the standard concentration of 4 mM dNTPs (Fig. S1 and Table 2). These results likely reflect that when the incoming dNTP can base pair with a templating nucleotide, 4 mM dNTP is saturating for the WT enzyme and nearly saturating for the mutant. Thus, under these conditions the F143A mutation selectively inhibits NTA activity.

### The F143A mutation decreases secondary template switches in RNA-seq

Template switching from an RNA template/DNA primer duplex similar to that in the previous experiments but with a 1-nt 3’-DNA overhang that is an equimolar mix of all four nucleotides (denoted N) is employed in a high-throughput RNA sequencing method called TGIRT-seq to initiate reverse transcription at the 3’ end of target RNAs with concomitant attachment of an RNA-seq adapter to the 5’ end of a cDNA (Fig. 7*A*; (34–36)). A second RNA-seq adapter is then added to the 3’ end of the completed cDNA by a single-stranded DNA ligation followed by minimal PCR amplification to generate the final TGIRT-seq library for sequencing (Fig. 7*A*). For this method, it is desirable to maximize the efficiency of the initial template switch from donor duplex with a pre-formed 1-nt 3’-DNA overhang used for RNA-seq adapter addition, while decreasing secondary template switches from the 3’ end of the completed cDNA, which result in chimeric reads that are typically discarded. The problem of multiple sequential template switches is most pronounced for TGIRT-seq of size-selected short RNAs, such as miRNAs (23, 36). Although secondary template switches in TGIRT-seq can be suppressed by a high salt concentration (450 mM NaCl), high salt also decreases the efficiency of template switching from the initial starter duplex (35), resulting in a lower yield of library product. A recently introduced modification of the TGIRT-seq method using a lower salt concentration (200 mM NaCl) for sequencing of human plasma RNAs resulted in < 5% soft clipped or discordant read pairs that could include secondary template switching (37), a manageable number but one that it would still be desirable to decrease further, as well as enable use of even lower salt concentrations to further increase the efficiency of the initial template-switching step.

**Figure 7.**
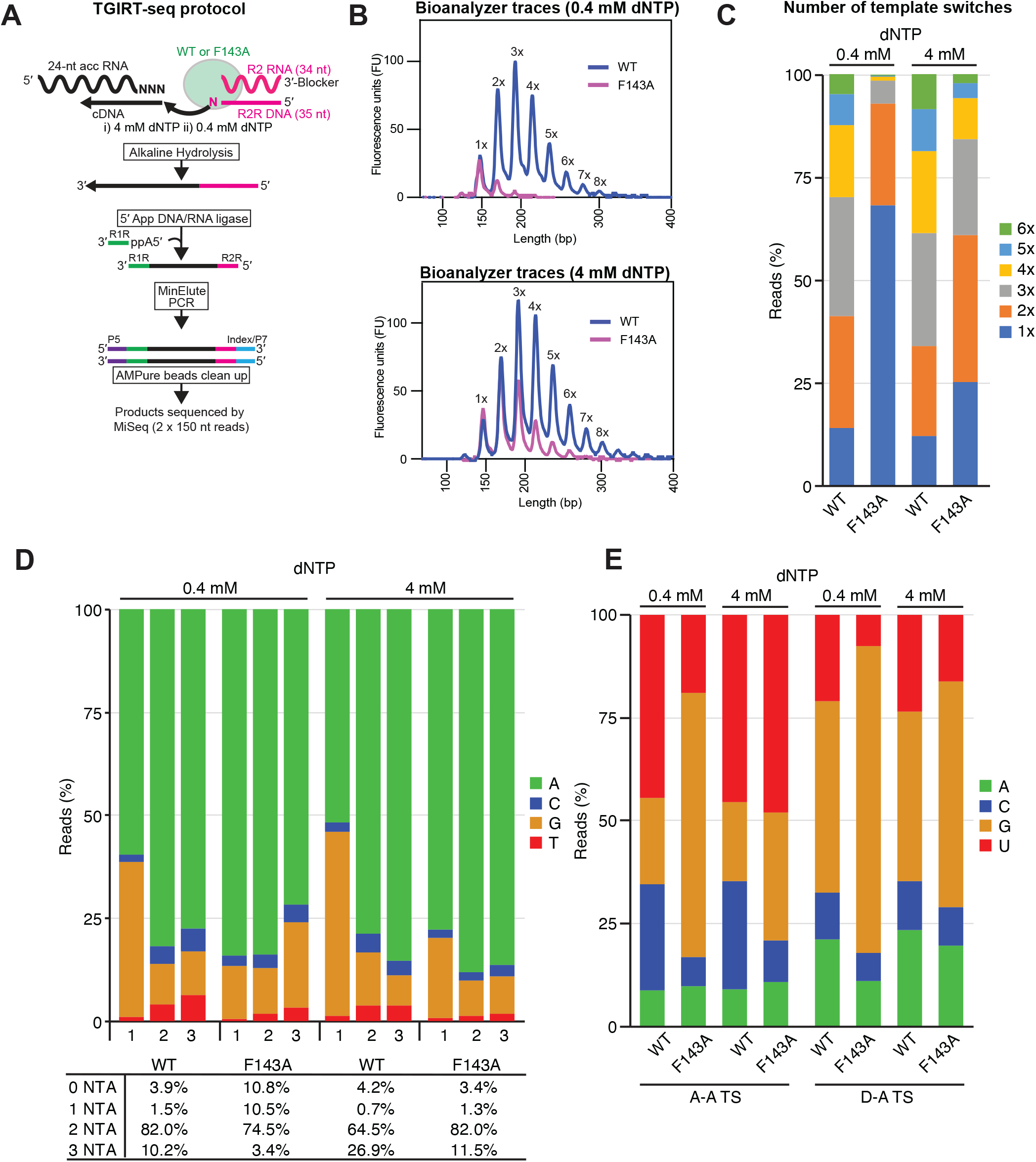
The F134A mutation decreases secondary template switches in TGIRT-seq. *A*, outline of the RNA-seq workflow. Template-switching reactions using WT or F143A GsI-IIC RTs were performed with an unlabeled starter duplex with a 1-nt 3’-DNA primer overhang that was an equimolar mixture of A, C, G, and T residues (denoted N) and a 24-nt acceptor RNA of which each of the last three nucleotides was an equimolar mixture of A, C, G, and U residues (denoted N). After a 30-min pre-incubation, reactions were initiated by adding dNTPs, incubated at 60 °C for 15 min, and terminated by adding NaOH and heating to 95 °C for 3 min. cDNA products were cleaned up by using a MinElute column (Qiagen) and then ligated to a 5′-adenylated R1R adapter using Thermostable 5′ App DNA/RNA Ligase (New England Biolabs). After another MinElute clean up, Illumina RNA-Seq capture sites (P5 and P7) and indices were added by PCR, and the resulting libraries were cleaned up by using AMPure XP beads (Beckman Coulter) prior to sequencing on an Illumina MiSeq v2 to obtain 150-nt paired-end reads. *B,* bioanalyzer (Agilent; High Sensitivity DNA chip) traces of TGIRT-seq libraries prepared via template switching at 0.4 mM and 4 mM dNTPs to the 24-nt acceptor RNA. The lengths indicated on the x axis are those of internal DNA size markers, which were omitted from the trace for clarity. *C*, stacked bar graphs show the percentages of reads containing a single template switch from the starter duplex (1x) or multiple template switches from the 5’ end of one RNA template to the 3′ end of another (2x to 6x) in TGIRT-seq datasets obtained with wild-type GsI-IIC RT and the F143A mutant at 0.4 or 4 mM dNTPs. *D*, stacked bar graphs showing the proportions of different nucleotides for the first, second, and third non-templated-nucleotide additions at the 3’ end of the final cDNA synthesized after the last template switch in TGIRT-seq datasets obtained with wild-type GsI-IIC RT and the F143A mutant at 0.4 and 4 mM dNTPs. The table below shows the percentages of 0, 1, 2, or 3 nt NTAs for each condition. *E*, stacked bar graphs showing nucleotide frequencies of the 3’-terminal nucleotide of the acceptor RNA in TGIRT-seq datasets obtained with wild-type GsI-IIC RT and the F143A mutant at 0.4 and 4 mM dNTPs for acceptor to acceptor (A-A) and starter duplex to acceptor (D-A) template switches.

To test the feasibility of using the F143A mutation to decrease the frequency of secondary template switches in TGIRT-seq, we constructed TGIRT-seq libraries with WT and F143A GsI-IIC RTs using the protocol outlined in Fig. 7*A* in reaction medium containing 200 mM NaCl under conditions designed to exacerbate the number of secondary template switches, including the use of a very short 24-nt acceptor RNA with three randomized 3’ nucleotides (denoted NNN) to enable base pairing of different nucleotides added by NTA to the 3’ end of completed cDNAs (Fig. 7*A*). Bioanalyzer traces of the final TGIRT-seq libraries after PCR amplification showed that F143A mutation substantially decreased the proportion of secondary template switches compared to the WT enzyme and that this decrease was more pronounced at 0.4 mM than at 4 mM dNTP (Fig. 7*B*), as expected from the experiments above comparing the efficiency of the reaction at subsaturating and near saturating dNTP concentrations for the mutant (Fig. 6).

Sequencing of the TGIRT-seq library on an Illumina MiSeq instrument showed that 68.5% of the reads obtained with the F143A mutant at the lower dNTP concentration were free of multiple template switches compared to only 13% for the WT enzyme at either dNTP concentration and 25% for the F143A mutant at the higher dNTP concentration (Fig. 7*C*). The F143A mutation did not increase the frequency of base substitutions or indels above the composite error rate for TGIRT-seq library preparation, oligonucleotide synthesis, PCR, and sequencing (Table S2). Because we used an acceptor RNA with randomized 3’ ends and because NTA still occurs to the 3’ end of the final cDNA synthesized after multiple template switches, the TGIRT-seq datasets could also be used to assess the effect of the F143A mutation on nucleotide preferences for both NTA and acceptor to acceptor (A-A) template switching. Notably, the F143A mutation led to a marked decrease in the proportion of G residues for the first NTA at both dNTP concentrations (Fig. 7D) and a concomitant decrease in secondary template-switching to acceptors with a complementary 3’ C residue (Fig. 7E). Collectively, these findings indicate that the F143A mutation in combination with lower dNTP concentrations may be useful for decreasing multiple template switches in TGIRT-seq. Additionally, because F143 corresponds to a conserved aromatic residue in the dNTP-binding site of all RTs, our findings suggest that mutations at this position may be generally useful for optimizing the number and composition of NTAs by retroviral and other RTs for SMART-seq and other RNA-seq applications.

### Reversible binding of acceptor RNAs within the template switching pockets and effect of RNA length

We hypothesized that the binding of the acceptor RNA within the template-switching pocket of GsI-IIC RT might be highly reversible, so that mismatched acceptors whose 3’ end is not complementary to the 1-nt DNA overhang could dissociate rapidly, while a matched acceptor with a complementary 3’ end might dissociate more slowly, enabling it to be used preferentially upon addition of dNTPs. To test this hypothesis, we probed whether the complex formed after binding an acceptor with a 3’ nucleotide complementary to the 1-nt DNA overhang is committed to extension or could exchange rapidly with an acceptor RNA from solution. In these experiments, we preincubated GsI-IIC RT with a saturating concentration of a 21-nt acceptor RNA and a donor template-primer duplex with a complementary 1-nt 3’-DNA overhang for 30 min to allow binding of the acceptor. We then initiated reverse transcription by adding dNTPs, with or without a 20-fold excess of a competitor 34-nt acceptor (Fig. 8*A*, top). To preclude sequence biases for binding within the template-switching pocket from influencing the experiment, the longer acceptor had the same 21-nt 3’ sequence as the shorter acceptor and differed only by the extra nucleotides at its 5’ end. Indeed, we found that template switching occurred predominantly to the competitor 34-nt acceptor (Fig. 8*A*, left lanes), indicating that despite the potential for base pairing with the 1-nt DNA overhang, the 21-nt acceptor binds reversibly, dissociating faster than dNTP binding and extension.

**Figure 8.**
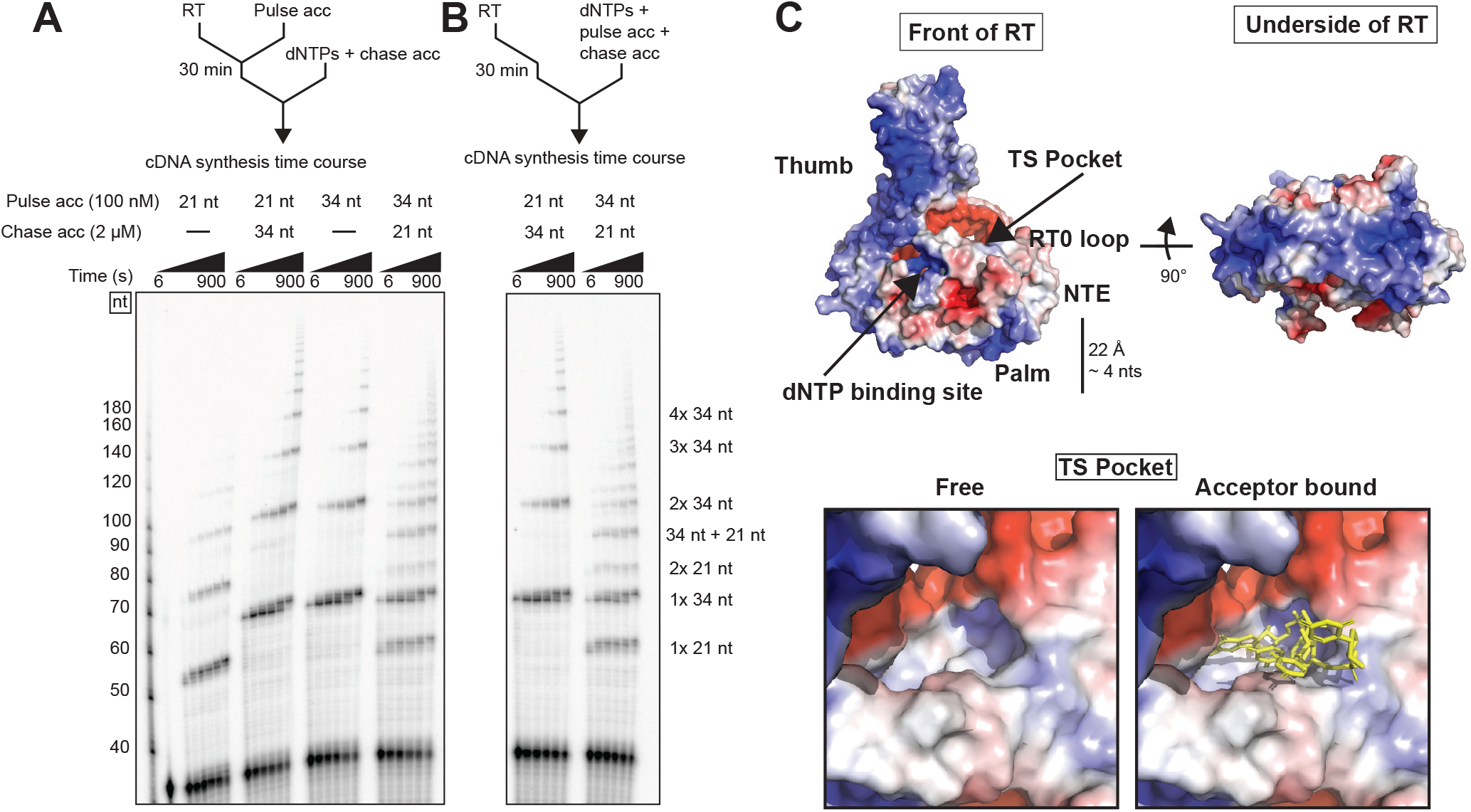
Template switching by GsI-IIC RT favors a longer acceptor RNA. *A*, schematic of the experiment. WT GsI-IIC RT (200 nM) in complex with a labeled 20 nM 1-nt 3’-DNA overhang starter duplex was incubated with 100 nM of a 21- or 34-nt acceptor RNA for 30 min, after which the reaction was started by adding 4 mM dNTPs and an excess (2 µM) of a reciprocal 34- or 21-nt acceptor RNA. *B*, WT GsI-IIC RT (200 nM) in complex with a labeled 20 nM 1-nt 3’-DNA overhang starter duplex was incubated for 30 min, after which the reaction was started by adding 4 mM total dNTPs, 100 nM of a 21- or 34-nt acceptor RNA and 2 µM of a 34-nt or 21-nt acceptor RNA. Reaction time courses were analyzed by electrophoresis in a denaturing 6% polyacrylamide. The numbers to the left of the gel indicate the lengths of size markers (5′-^32^P-labeled single-stranded DNA ladder; ss20 DNA Ladder, Simplex Sciences) run in a parallel lane of the same gel. *C*, electrostatic potential surface representation of GsI-IIC RT created using the APBS Electrostatics plugin for Pymol. Electropositive regions are blue, and electronegative regions are red. In the front view (top left), the entrance of the TS pocket is indicated by an arrow, and a scale bar is included to give a sense of the distance that could be reached from the 5’ end of an acceptor RNA extending outside the template-switching pocket. Close up views of the template-switching pocket without and with the bound 5-nt acceptor RNA in the crystal structure (yellow stick) are shown at the bottom.

Surprisingly, in a reciprocal experiment, in which GsI-IIC RT was preincubated with the 34-nt acceptor RNA and the donor duplex for 30 min before adding dNTPs with a 20-fold molar excess of 21-nt acceptor, only ∼50% of the template-switching product switched to the shorter 21-nt acceptor (Fig. 8*A*, right lanes). Although this result could have indicated that the complex formed with the 34-nt acceptor is equally likely to dissociate or to go on to chain extension, it was also possible that the 34-nt acceptor RNA binds preferentially from solution, even though both acceptors have identical 3’-end sequences.

To distinguish these possibilities, we performed an additional experiment in which both the longer and shorter acceptors were added together in different ratios along with dNTPs to initiate reverse transcription (Fig. 8*B*). This experiment showed that when the 34-nt acceptor was present in excess from the beginning, it was used exclusively for template switching, whereas when the 21-nt acceptor was present in excess from the beginning, the 34-nt was still present in > 50% of the template-switching products. This preferential binding of the longer acceptor RNA indicates that its 5’ end can interact with additional sites on the RT outside of the template-switching pocket, likely patches of basic residues on the outer surface of the protein that are used for binding group II intron RNAs for RNA splicing and retrohoming (24) (Fig. 8*C*). These findings account for a previously observed size bias against RNAs < 60 nt in heterogeneously sized RNA preparations in TGIRT-seq using GsI-IIC RT (37, 38) and suggest approaches for mitigating this bias by mutating basic surface residues that are not required for reverse transcription.

## Discussion

Here, we determined a crystal structure of a group II intron RT template-switching complex with synthetic oligonucleotides representing an acceptor RNA and a donor RNA template/cDNA duplex after NTA to form a complementary 1-nt 3’-DNA overhang. The structure showed that the 3’ end of the acceptor RNA binds in a pocket formed by the NTE and fingertips loop with its 3’ nucleotide base paired to the 1-nt DNA overhang and its penultimate nucleotide base paired to an incoming dNTP at the RT active site. This template-switching pocket is ordinarily occupied by upstream regions of a continuous RNA template but would become vacant when the RT reaches the 5’ end of an RNA template or dissociates from a broken or damaged template, enabling template switching to the 3’ end of a new acceptor RNA template for the continuation of cDNA synthesis.

Globally, the structure of the GsI-IIC RT template-switching complex is similar to that of GsI-IIC RT bound to a continuous RNA template/DNA primer duplex, the major differences detectable at this resolution being the lack of a phosphate in the gap between the 3’ end of the acceptor and 5’ end of the donor templates and a small shift in the position of N23, possibly reflecting the lack of that phosphate. The NTE, which forms part of the template-switching pocket, is a common feature of non-LTR-retroelement RTs and viral RdRPs but is lacking and presumed to have been lost from retroviral RTs. A critical part of the NTE for template switching is the RT0 loop, a structured polypeptide linker which in GsI-IIC RT connects the two bent *α*-helices of the NTE and forms a lid over the RNA template backbone (Fig. 2 and 4). The RT0 loop sequence is conserved in group II intron RTs and partially conserved in non-LTR-retroelement RTs (Fig. 1C). In GsI-IIC RT, conserved small amino acids at the N-terminus of the RT0 loop (G25 and A26) allow close approach and interaction of this portion of the lid with the RNA acceptor phosphate backbone. Most of the remaining conserved amino acids in the loop are hydrophobic residues involved in packing interactions with the fingers that anchor the middle and C-terminal end of the structured loop. In high-resolution crystal structures of two closely related apo-group II intron RTs (29), the RT0 loop adopted the same conformation utilizing the same conserved interactions. Further, cryo-EM structures of the more distantly related Ll.LtrB and TeI4h group II intron RTs in complexes with group II intron RNA support a similar location for the lid and a packed RT0 loop in the absence of reverse transcription substrates (28, 30). Collectively, these findings indicate that the structured RT0 loop conformation as seen in GsI-IIC RT is not dependent upon the presence of a nucleic acid substrate in the template-switching pocket. Additionally, the RT0 loop has the same configuration in the template-switching structure and the continuous structure, suggesting that it may not undergo large conformational changes during template switching.

Because the 3’ end of the RNA acceptor is bound and held in place primarily by polypeptide backbone interactions with the RT0 loop lid, we sought to disrupt the structure of the lid to test whether acceptor binding would be affected. We found that the mutation 23-31/4G. in which residues 23-31 were replaced by four glycines, resulting in a shorter loop and ablating a critical anchoring interaction of the loop with the fingers, strongly inhibited template-switching activity but had no detectable effect on primer-extension activity, indicating that maintaining the proper conformation of the lid is critical for acceptor RNA binding. The binding of the lid to the RNA acceptor may constrain mobility at its 3’ end and facilitate its alignment for template switching. After acceptor RNA binding, the shortened, glycine-rich lid in the 23-31/4G mutant adopts a conformation that does not impede reverse transcription.

Most of the remainder of the binding surface for the acceptor RNA in the template-switching pocket is formed by the fingertips loop, which is a conserved β-hairpin that is present in all RTs and harbors the two essential basic catalytic residues (K69 and R75 in GsI-IIC RT) on the surface that faces the polymerase active site (31). In known structures of RTs that contain a 5’-RNA overhang, the opposite face of the fingertips loop binds to the single-stranded RNA template region that awaits copying by using a group of hydrophobic residues projecting outward from the *β*-hairpin (39, 40). In the template-switching structure, the acceptor RNA binds to this same surface of the fingertips loop via a series of aliphatic residues that are conserved in other group II intron and non-LTR-retroelement RTs (Fig. 1*C*). We found that mutations in L77 and I79 strongly inhibited template-switching activity, while only slightly slowing primer extension. These residues comprise the primary binding surface for the T-1 and T-2 nucleotides of the acceptor RNA and collectively contact most of the fingertips loop-facing surface of the ribose rings and bases of these nucleotides. The finding that mutations in these residues strongly inhibited template switching but not primer extension indicates that they make a proportionally greater contribution to binding an acceptor RNA for template switching than for binding the upstream region of a continuous RNA template, which is covalently attached to the polymerized duplex.

In the template-switching structure, the RNA acceptor binds to a conformation of the fingertips loop that is engaged with the dNTP via the catalytic basic residues and poised for reverse transcription. In group II intron RT structures that do not contain substrate, the fingertips loop is observed in a more open conformation, although these structures are also influenced by the presence of the intron RNA (28, 30) or crystal packing interactions (29). Presumably, though, the fingertips loop must be somewhat dynamic to allow binding of a new dNTP after each polymerization cycle. In the open conformation, the catalytic basic residues are out of reach of the dNTP, and a portion of the RNA acceptor binding site would be disrupted. Our structure suggests that binding of dNTP and RNA acceptor could work in concert to promote reverse transcription during a template switch by stabilizing the closed conformation of the fingertips loop.

We found previously that efficient template switching is dependent upon base pairing of the 3’ nucleotide of the acceptor with a 1-nt 3’-DNA overhang representing NTA of a single nucleotide to the 3’ end of the cDNA and that longer overhangs resulting from multiple NTAs were progressively less efficient, possibly reflecting the difficulty of matching two base pairs within the narrow confines of the template-switching pocket prior to initiating cDNA synthesis (27). Here, we identified a mutation in a phenylalanine residue (F143A), which strongly inhibited NTA, while retaining relatively high primer-extension activity (Fig. 6 and Fig. S1). F143 corresponds to a conserved aromatic residue (F or Y) that is present in in all RTs and forms a pi-stacking interaction with the deoxyribose of the incoming dNTP, while also serving to exclude rNTPs from the active site via a steric clash with the 2’ OH of the rNTP ribose (24,33,39). During NTA, when the incoming dNTP is not held in place by base pairing with a templating RNA nucleotide, the F143A mutant displays substantially decreased activity even at high dNTP concentrations, when compared to primer extension for which dNTP concentrations remain close to saturating (compare Fig. S1 to Fig. 6C and Fig. S5A). Our finding that this mutation inhibited template-switching from the 5’ end of a reverse transcribed template or a blunt-end donor RNA/DNA duplex (Fig. 6C and D), but not from a donor RNA/DNA duplex with a pre-formed 1-nt DNA overhang, confirms the necessity of NTA after the completion of cDNA synthesis for template switching to a new acceptor.

Previous studies showed that the specificity of template-switching by GsI-IIC RT is dictated largely by base pairing of the 3’ end of the acceptor to the 1-nt 3’-DNA overhang of the donor, with no sequence biases for other positions at the 3’ end of the acceptor RNA (23,34,36). The template-switching structure indicates that this lack of sequence bias reflects that binding of the acceptor within the template-switching pocket is dependent largely upon either hydrophobic interactions or interactions of the protein with the phosphate backbone or ribose 2’ OH, with no H-bond interactions to specific bases. This weak binding of the acceptor enables its rapid dissociation in the absence of a base pair with a complementary 1-nt 3’-DNA primer overhang added by NTA or provided by a starter duplex for RNA-seq.

Remarkably, despite being dictated by only a single base pair, the template-switching reaction of GsI-IIC RT has high specificity, with RNA-seq showing 97.5-99.7% precise junctions depending upon the base pair (27). Our structure and kinetic analysis suggest that this high specificity reflects that after base pairing of the 3’ nucleotide of the acceptor to the 1-nt DNA overhang, a second base-pairing interaction between the templating RNA base and the incoming dNTP accompanied by a conformational change involving the fingertips loop results in initiation of reverse transcription, making the reaction with the matched acceptor irreversible.

Several of our findings are relevant to the use of group II intron RTs for RNA-seq. First, the F143A mutation, which inhibits NTA and secondary template switches (Fig. 6 and Fig. 7), provides a new tool for decreasing artifactual fusion reads, making it easier to identify *bona fide* hybrid transcripts, such as those resulting from chromosome rearrangements in cancer and other diseases. The use of this mutation would decrease reliance on high salt concentrations, which have been used previously to decrease secondary template switches but also decrease the efficiency of the initial template switch used for RNA-seq adapter addition from a 1-nt 3’ overhang duplex (35). We also found that the F143A mutation changes nucleotide preferences for NTA and concomitantly changes nucleotide preferences for template switching from the 5’ end of one template to the 3’ end of another, which are dependent upon base pairing of the nucleotides added by NTA to the 3’ end of the new acceptor (Fig. 7, *D* and *E*). As F143 corresponds to a conserved aromatic found in all RTs (33), engineering of this residue to ablate, change, or optimize the pi-stacking interaction to modulate NTA might also be possible for retroviral and other RTs employed in RNA-seq methods such SMART-seq. Finally, our finding that longer RNAs out compete shorter RNAs with identical 3’ ends indicates that the efficiency of template switching can be influenced by the binding of upstream regions of longer acceptor RNAs to secondary sites outside the template-switching pocket and accounts for the previously observed TGIRT-III bias against miRNA-sized RNAs in heterogeneously sized RNA preparations (36). These secondary binding sites likely include patches of basic residues on the surface of group II intron RTs that are used for binding group II intron RNAs for RNA splicing and reverse splicing in retrohoming. These basic patches are not required for reverse transcription and thus should be dispensable and relatively easy to remove for RNA-seq applications (41, 42).

Although all non-LTR-retroelements RTs have an NTE, both its structure and the degree of conservation of the RT0 loop vary in different enzymes. In group II intron RTs, the NTE appears to be predominantly *α*-helical, but the arrangement of the helices varies in known group II intron RT structures (28, 30). The hypothetical RT0 loops of insect R2 and human LINE-1 related RTs share only a glycine residue, which lies outside of the lid sequence, and an aspartate residue (homologous to D30 in GsI-IIC RT), which appeared largely dispensable for template switching. Despite this lack of sequence conservation, mutating the RT0 loop in the *Bombyx mori* R2 element RT abrogated template switching activity (43), similar to our findings for GsI-IIC RT. In contrast to the RT0 loop sequence, the hydrophobic, aliphatic residues that contribute to acceptor RNA binding in the fingertips region of the template-switching pocket of GsI-IIC RT are well conserved in non-LTR-retroelement RTs (Fig. 1*C*). Collectively, these findings together with other previous biochemical analyses of template switching by the R2 element RT (44, 45) suggest that group II intron and non-LTR-retroelement RTs use fundamentally similar template-switching mechanisms but that the structure of the NTE and template-switching pocket can diverge, possibly to accommodate the life cycle of different retroelements.

An example of such accommodation is provided by *Neurospora* spp. mitochondrial retroplasmids, which encode RTs that have a predominantly helical NTE domain and an RT0 loop region with no recognizable sequence similarity to group II intron RTs (46). These retroplasmid RTs use template switching to a 3’ CCA sequence to generate concatemeric DNAs that are resolved by intramolecular DNA recombination to generate circular plasmid DNAs (47, 48). They also template switch to the 3’ end of tRNAs to generate chimeric plasmids for which the integrated tRNA sequence confers a replicative advantage when integrated into plasmid DNA (49). RT recognition of the 3’ CCA could occur within the template-switching pocket, possibly involving the RT0 loop, which contains multiple aromatic Phe, Tyr, and Trp residues that could mediate recognition of specific nucleotides (46). In addition to the 3’ CCA, the retroplasmid RT also binds the acceptor stem cognate of a 3’ tRNA-like secondary structure of the plasmid transcript (47), possibly using surface residues outside the template-switching pocket, as suggested for the binding of longer oligonucleotides with longer 5’ ends by GsI-IIC RT (Fig. 8*B*).

We found that group II intron RTs utilize the same binding surface for both upstream region of continuous RNA templates during primer extension and acceptor binding during template switching, with the acceptor sandwiched between the NTE/RT0 loop on one side and the fingertips loop on the other side (Fig. 9*A*). Retroviral and LTR-retrotransposon RTs, which lack an NTE with an RT0 loop, perform analogous template switching termed clamping, yet typically require at least two base pairs of primer-acceptor overlap for efficiency instead of a single base pair as for group II intron RTs (19, 20). Based on our findings for GsI-IIC RT, modeling suggests that the acceptor-binding surface for retroviral RTs consists of residues in the palm near the primer-acceptor overlap and hydrophobic residues in the fingertips loop in homologous positions to those found in GsI-IIC RT (Fig. 9*B*) (19, 20). The lack of any protein structure corresponding to the NTE/RT0 loop to sandwich the RNA acceptor may explain the necessity of at least one additional base pair and a larger binding surface on the palm for efficient clamping by retroviral RTs compared to group II intron RTs. Retroviral RTs can also template switch via strand transfer, but this process is more akin to primer extension, as the donor template is degraded by RNase H and the remaining cDNA base pairs to a complementary template (18).

**Figure 9.**
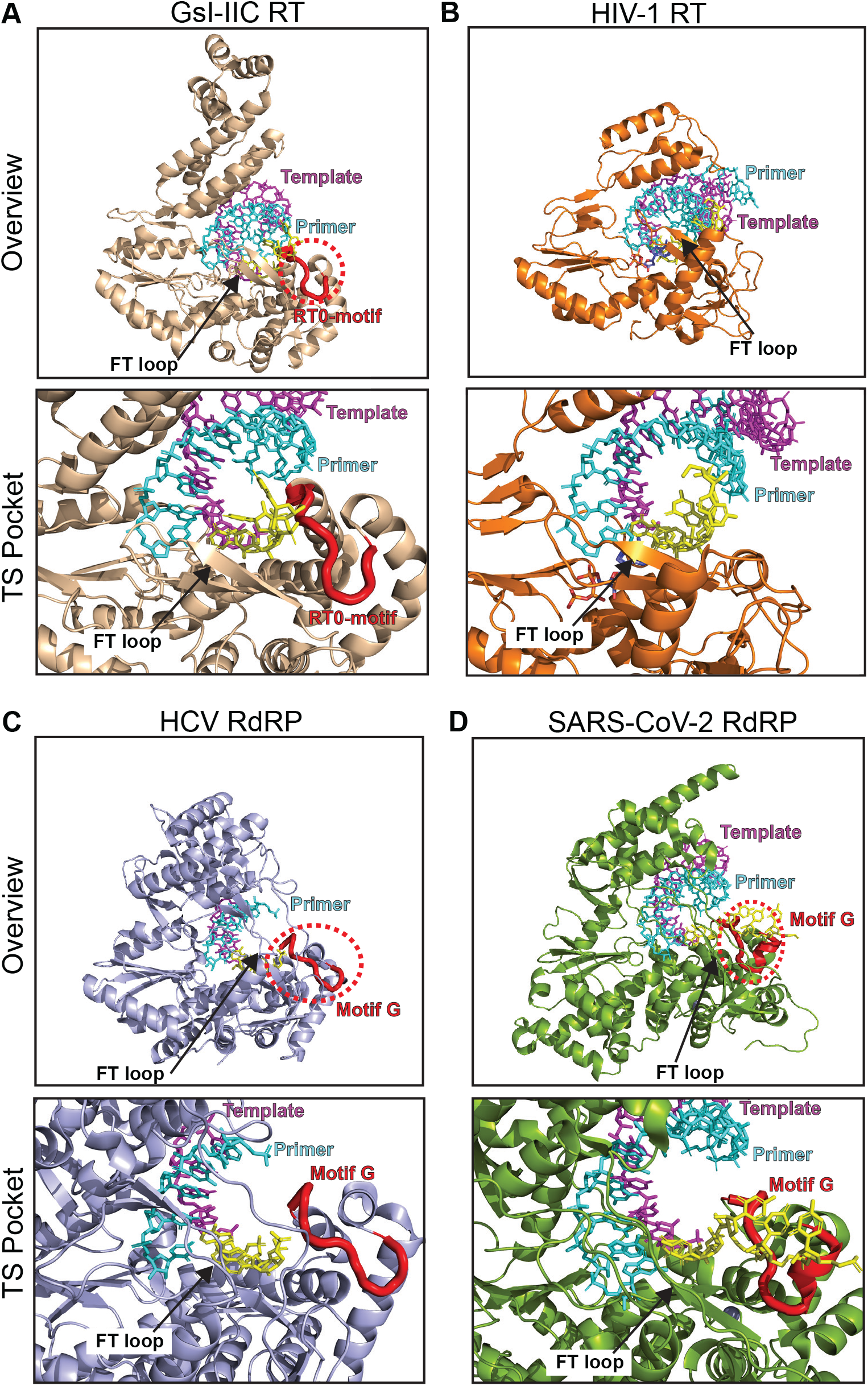
Comparison of upstream template RNA-binding regions that comprise the template-switching pocket of GsI-IIC RT compared to the same regions of HIV-1 RT and viral RdRPs. *A*, GsI-IIC RT (PDB: 6AR1 (24), tan); *B,* HIV-1 RT (PDB: 4PQU (39), amino acids 1 to 312 showing the RT and thumb domains, orange); *C,* HCV RdRP (PDB: 4WTA (56), light blue); *D*, SARS-CoV-2 RdRP (PDB: 72CK (57) with the non-homologous N-terminal amino acids 1-356 removed, light green). For each enzyme, cartoon representations are shown as overviews (top) or zoomed in on the putative template-switching pocket region (bottom). Primer strands are cyan (stick representation) and RNA strands are purple (stick representation) in all three proteins. The RT0 loop and the RT0 loop cognate ‘motif G’ in RdRPs are highlighted and shown in red carton within a dotted red circle. The fingertips (FT) loop is indicated by an arrow. Structural alignments were done using Coot (64).

Finally, viral RdRPs, which are thought to be evolutionary ancestors of RTs, can template switch by an end-to-end mechanism analogous to clamping as well as by strand transfers that produce physiologically relevant subgenomic RNAs and enable recombination between viral RNA templates (50–55). Structures of well-studied RdRPs, including HCV RdRP (4WTA (56) and SARS-Cov-2 RdRP (72CK (57) (Fig. 9, *C* and *D*; see also PDB:2E9T (58), PDB:5TSN (59), and PDB:1S48 (60)) show a putative TS pocket composed of an RT0 loop cognate (termed motif G in RdRPs) and the fingertips loop on either side. The template-switching pocket homology between RdRPs and group II intron RTs suggests a common mechanism for binding the acceptor RNA template. Furthermore, small numbers of extra nucleotides at TS junctions for some viruses suggest a role for NTA similar to the mechanisms of retroviral and group II intron RT clamping (27, 52). However, RdRPs lack an aromatic residue corresponding to the NTA-facilitating F143 in GsI-IIC RT, which would sterically clash with the 2’ OH on the ribose ring of rNTPs, and instead use hydrogen bonds to the 2’ OH to impart selectivity for rNTPs (33). Additionally, in HCV RdRP, the mutation D225A in a residue that is part of a conserved hydrogen-bonding network that stabilizes the ASG loop, inhibits template-switching but not primer extension or NTA, possibly by affecting interaction of the ASG loop with the acceptor RNA (54). As with viral RdRPs, group II intron RTs typically possess homologs of both D225 (D144 in GsI-IIC RT) and the ASG loop (PQG loop in GsI-IIC RT with G192 contacting T-1 of the acceptor; Fig. 1B), suggesting future avenues for investigating the effect of mutations in these residues on template-switching activity in these polymerase families.

## Acknowledgements

High-throughput sequencing was done by the Genomic Sequencing and Analysis Facility at the University of Texas at Austin. The Center for Biomedical Research Support (CBRS) at the University of Texas at Austin provided high-performance computing resources.

## Author contributions

A.M. Lambowitz initiated the project. A.M. Lambowitz, A.M. Lentzsch and R.R. devised the experiments. A.M .Lentzsch conducted all the experiments. J.Y. analyzed the RNA-seq data. A.M. Lentzsch purified and crystallized the GsI-IIC RT protein. A.M. Lentzsch and J.S. solved the crystal structure. All authors wrote and edited the article.

## Funding and additional information

This work was supported by National Institutes of Health Grants R35 GM136216 (to A.M. Lambowitz) and R35 GM131777 (to R.R.). Beamline 5.0.1 of the Advanced Light Source, a U.S. DOE Office of Science User Facility under Contract No. DE-AC02-05CH11231, is supported in part by the ALS-ENABLE program funded by the National Institutes of Health, National Institute of General Medical Sciences, grant P30 GM124169-01. The content is solely the responsibility of the authors and does not necessarily represent the official views of the National Institutes of Health.

## Experimental procedures

### DNA and RNA oligonucleotides

The DNA and RNA oligonucleotides used in this work are listed in Table S1. Most were purchased in RNase-free, HPLC-purified form from Integrated DNA Technologies. For biochemical assays, the R2R and primer extension (PE) DNA primers (100 pmol) were labeled with [γ-^32^P]ATP (125 pmol; 6,000 Ci/mmol; 150 μCi/ μl; PerkinElmer Life Sciences) by incubating the DNA with T4 polynucleotide kinase (10 units; New England Biolabs) in the reaction medium provided by the supplier for 30 min at 37 °C. A typical 10-μl reaction was then diluted to 40 μl with double-distilled H_2_O and extracted with an equal volume of phenol/chloroform/isoamyl alcohol (25:24:1). Unincorporated nucleotides were removed from the aqueous phase by using a P-30 Microspin column (RNase Free; Bio-Rad), and the oligonucleotide concentration was measured by using a Qubit ssDNA assay kit (Thermo Fisher Scientific). The concentration of unlabeled oligonucleotides was determined by spectrophotometry using a NanoDrop 1000 (Thermo Fisher Scientific).

### Expression and purification of GsI-IIC RT protein used in crystallography

GsI-IIC RT with a C-terminal 8x His tag was expressed from plasmid pET-14b GsI-IIC RT C-term 8xHis and purified by essentially the same methods described previously (24). A freshly transformed colony of *Escherichia coli* BL21-CodonPLus (DE3)-RIPL cells (Agilent) was inoculated into 100 ml of Luria-Bertani (LB) medium containing ampicillin (50 μg/ml) and chloramphenicol (25 μg/ml) in a 250-ml Erlenmeyer flask and grown overnight with shaking at 37 °C. 50 ml of this starter culture was then added to 1 liter of LB medium containing ampicillin (50 μg/ml) in a 4-liter Erlenmeyer flasks and grown at 37 °C to an O.D._600_ of 0.7-0.8, at which time protein expression was induced by adding 1 mM isopropyl 1-thio-β-D-galactopyranoside (IPTG) and incubating at 37 °C for 2 h. The cells were pelleted by centrifugation and stored at –80 °C overnight. After thawing on ice, the cells were lysed by sonication in 40 ml of buffer containing 100 mM NaCl, 20 mM Tris-HCl pH 8.5, 10% glycerol, 5 mM imidazole, 0.1% β-mercaptoethanol, 0.2 mM phenylmethylsulfonyl fluoride (PMSF), and 1 cOmplete EDTA-free protease inhibitor cocktail tablet (Roche), and the lysate was clarified by centrifugation at 30,000 x *g* for 60 min at 4 °C in a JA25.50 rotor in an Avanti J-E centrifuge (Beckman). The clarified lysate was then added to 10 mL of Ni-NTA Agarose beads (Invitrogen) and incubated with rotation at 4 °C for 2 h. The beads were washed under gravity flow first with 250-ml Wash Buffer A (20 mM Tris-HCl pH 8.5, 100 mM NaCl, 10% glycerol, 5 mM imidazole and 0.1% β-mercaptoethanol) and then with 150-ml Wash Buffer B (20 ml Tris-HCl pH 8.5, 100 mM NaCl, 10% glycerol, 50 mM imidazole and 0.1% β-mercaptoethanol). The protein was then eluted by washing 5 times with 5 ml of Elution buffer (20 mM Tris-HCl pH 8.5, 100 mM NaCl, 10% glycerol, 250 mM imidazole and 0.1% β-mercaptoethanol). Fractions were analyzed by SDS-PAGE, and those containing GsI-IIC RT were pooled and loaded onto a 5-ml Heparin HP column (GE Healthcare) pre-equilibrated in Heparin Buffer A (20 mM Tris-HCl pH 8.5, 100 mM NaCl, 10% glycerol, and 0.1% β-mercaptoethanol) at a flow rate of 1 ml/min. The column was washed with 5 column volumes (CV) of Heparin Buffer A and eluted with a 10 CV linear gradient of Heparin Buffer A to 50% Heparin Buffer B (2 M NaCl, 20 mM Tris-HCl pH 8.5, 10% glycerol, and 0.1% β-mercaptoethanol), after which a final 5 CV of 100% Heparin Buffer B was applied. Fractions containing GsI-IIC RT identified by SDS-PAGE were pooled and concentrated by incubating in 65% saturating ammonium sulfate and centrifuging the precipitated protein at 20,000 x *g* for 2 h at 4 °C. The resulting protein pellet was resuspended in crystallization buffer (500 mM NaCl, 20 mM Tris-HCl pH 8.5, 10% glycerol, and 5 mM dithiothreitol (DTT) to a final concentration of ∼ 2 mg/ml.

### Crystallization

For crystallization, an RNA/DNA starter duplex was formed by combining the single-stranded RNA template and DNA primer oligonucleotides, the latter ending with 3’ ddCTP, at a 1:1 molar ratio, heating to 82 °C for 2 min, and then slowly cooling to room temperature. The annealed starter duplex was then combined with GsI-IIC RT and a 5′-UUUUG acceptor RNA oligonucleotide at a molar ratio of 1:1.2:2 (protein : starter duplex : acceptor RNA). The crystallization buffer also contained 1 mM dATP and 10 mM MgCl_2_. This mixture was incubated on ice for 30 min and centrifuged at 20,000 x *g* for 30 min at 4 °C. Crystals were grown by the hanging drop vapor diffusion method, with drops containing 0.5 μL of GsI-IIC RT/duplex/acceptor and 0.5 μL of well solution containing 0.1 M HEPES pH 7.0 and 1.3 M sodium citrate tribasic dihydrate. Crystals grew as characteristic thin plates over the course of 1 to 2 weeks. Crystals were harvested with a cryoloop (Hampton Research) and immersed briefly in Al’s oil (Hampton Research) before flash freezing in liquid nitrogen.

### Data collection and processing

Diffraction data were collected at 100K at the Advanced Light Source (ALS) on beamline 5.0.1. Images were integrated using the XDS package (61) and scaled with Aimless (62). Molecular replacement followed by rigid body refinement was carried out using a GsI-IIC RT model (PDB: 6AR3) with the dATP and the RNA acceptor removed. Refinement was carried out in Refmac5 with the twin law h+2*l, -k, -l, twin fraction = 0.5 applied (63). The R_free_ set was chosen to match that of 6AR3. Data collection and refinement parameters are reported in Table 1. The monomer composed of chains A (protein), B (DNA primer), C (RNA donor) and G (RNA Acceptor) was used throughout this work for depictions and structural analysis due to better density than the other monomer.

### Expression and purification of GsI-IIC RT proteins used in biochemical assays

GsI-IIC RT with an N-terminal maltose-binding protein tag to keep the protein soluble and stable in the absence of bound nucleic acids was expressed from plasmid pMRF–GsI-IIC RT (23). A freshly transformed colony of *E. coli* Rosetta 2 (DE3) cells (EMD Millipore) was inoculated into 1 liter of LB medium containing ampicillin (50 μg/ml) and chloramphenicol (25 μg/ml) in a 4-liter Erlenmeyer flask and grown overnight with shaking at 37 °C. 50 ml of the starter culture was then added to 1 liter of LB medium containing ampicillin (50 μg/ml) in one to six 4-liter Erlenmeyer flasks and grown at 37 °C to an O.D._600_ of 0.6-0.7, at which time protein expression was induced by adding 1 mM IPTG and incubating overnight with shaking at 19 °C. The cells were pelleted by centrifugation and stored at -80 °C overnight. After thawing on ice, the cells were lysed by sonication in 500 mM NaCl, 20 mM Tris-HCl pH 7.5, 20% glycerol, 0.2 mM PMSF (Roche Applied Science), 1 cOmplete EDTA-free protease inhibitor cocktail tablet (Roche), after which the lysate was clarified by centrifugation at 30,000 x *g* for 60 min at 4 °C in a JA 25.50 rotor in an Avanti J-E centrifuge (Beckman). Nucleic acids in the clarified lysate were precipitated by slowly adding polyethyleneimine with constant stirring in an ice bath to a final concentration of 0.4%, and then centrifuging at 30,000 x *g* for 25 min at 4 °C in a JA 25.50 rotor (Beckman). GsI-IIC RT and other cellular proteins were then precipitated from the supernatant with 60% saturating ammonium sulfate, pelleted at 30,000 x *g* for 25 min at 4 °C in a JA 25.50 rotor (Beckman), and resuspended in 25 ml of A1 buffer (300 mM NaCl, 25 mM Tris-HCl pH 7.5, 10% glycerol). The protein mixture was then purified through two tandem 5-ml MBPTrap HP columns (GE Healthcare). After loading, the tandem column was washed with 5 CVs of A1 buffer, and the maltose-binding protein–tagged GsI-IIC RT was eluted with 10 CVs of 500 mM NaCl, 25 mM Tris-HCl pH 7.5, 10% glycerol containing 10 mM maltose. The final fractions containing the RT were diluted to 200 mM NaCl, 20 mM Tris-HCl pH 7.5, 10% glycerol, loaded onto a 5-ml HiTrap Heparin HP column (GE Healthcare), and eluted with a 12-CV gradient from buffer A1 to A2 (1.5 M NaCl, 25 mM Tris-HCl pH 7.5, 10% glycerol) taking 0.5-ml fractions. Fractions containing GsI-IIC RT were identified by SDS-PAGE, pooled, and concentrated using a 30K centrifugation filter (Amicon). The concentrated protein was then dialyzed into 500 mM NaCl, 20 mM Tris-HCl pH 7.5, 50% glycerol, and aliquots were flash-frozen using liquid nitrogen and stored at –80 °C. Protein concentrations were determined by the Pierce BCA Protein Assay Kit (Thermo Fisher Scientific).

### Template-switching and non-templated addition assays

Template-switching reactions with GsI-IIC RT were carried out by using the previously described protocol with minor modifications (23). The RNA template/DNA primer starter duplex was based on that used for 3′-adapter addition in TGIRT-seq (34) and consists of a 34-nt RNA oligonucleotide containing an Illumina R2 sequence (R2 RNA) with a 3’-blocking group (3SpC3, Integrated DNA Technologies; Table S1) annealed to a complementary DNA primer (R2R) that leaves either a blunt end or a single nucleotide 3′-DNA overhang end (Table S1). The oligonucleotides were mixed in 1x TE at a ratio of 1:1.2, annealed by heating to 82 °C for 2 min, and then slowly cooled to room temperature to yield a donor duplex concentration of 20 nM. Unless specified otherwise, reactions were done with 200 nM GsI-IIC RT, 20 nM R2 RNA/R2R DNA starter duplex, and various concentrations of acceptor oligonucleotide in 25 μl of reaction medium containing 200 mM NaCl, 5 mM MgCl_2_, 20 mM Tris-HCl pH 7.5, 5 mM fresh DTT, and an equimolar mix of 1 mM each of dATP, dCTP, dGTP, and dTTP (Promega) to give a total dNTP concentration of 4 mM. Reactions were set up with all components except dNTPs, preincubated for 30 min at room temperature, and initiated by adding 1 μl of 25 mM dNTPs (an equimolar mix of 25 mM dATP, dCTP, dGTP, and dTTP). Reactions were then incubated at 60 °C for times indicated in the figure legends and stopped by adding 2.5-μl portions to 7.5 μl of 0.25 M EDTA. The products were further processed by adding 0.5 μl of 5 N NaOH and heating to 95 °C for 3 min to degrade RNA and remove tightly bound GsI-IIC RT, followed by a cooling to room temperature and neutralization with 0.5 μl of 5 N HCl. After adding formamide loading dye (5 μl; 95% formamide, 0.025% xylene cyanol, 0.025% bromophenol blue, 10 mM Tris-HCl pH 7.5, 6.25 mM EDTA), the products were denatured by heating to 99 °C for 10 min and placed on ice prior to electrophoresis in a denaturing 6% polyacrylamide gel (7 M urea, 89 mM Tris borate, and 2 mM EDTA, pH 8) at 65 watts for 1.5–2 h. A 5′-labeled, ssDNA ladder (10–200 nts: ss20 DNA Ladder, Simplex Sciences) was run in a parallel lane in all experiments. The gels were dried, exposed to an Imaging screen-K (Bio-Rad), and scanned using a Typhoon FLA 9500 phosphorimager (GE Healthcare). NTA reactions were done as described for template-switching reactions but in the absence of acceptor oligonucleotide.

### Primer-extension assays

Primer-extension reactions with GsI-IIC RT were carried out by using a 50-nt, 5′-labeled DNA primer (PE primer) annealed near the 3′ end of a 1.1-kb *in vitro*-transcribed RNA. The transcript was generated by T3 runoff transcription (T3 MEGAscript kit, Thermo Fisher Scientific) of pBluescript KS (+) (Agilent) linearized using XmnI (New England Biolabs) and cleaned up using a MEGAclear kit (Thermo Fisher Scientific). The labeled DNA primer was mixed with the RNA template at a ratio of 1:1.2 and annealed by heating to 82 °C for 2 min followed by slowly cooling to room temperature to a yield a final duplex concentration of 250 nM. GsI-IIC RT (200 nM) was pre-incubated with 20 nM of the annealed template–primer in 25 μl of reaction medium containing 200 mM NaCl, 5 mM MgCl_2_, 20 mM Tris-HCl pH 7.5, 5 mM DTT for 30 min at room temperature, and reverse transcription was initiated by adding 1 μl of the 25 mM dNTP mix to give a final dNTP concentration of 1 mM for each dNTP. After incubating at 60 °C for times indicated in the Figure Legends, the reaction was terminated, processed, and analyzed by electrophoresis in a denaturing 6% polyacrylamide gel, as described above for template-switching reactions.

### Analysis of kinetics experiments

Phosphorimager scans of reaction time courses were quantified with ImageQuant TL 8.1 (GE Healthcare) by generating rectangular boxes around the reaction products and unextended primer. For template-switching reactions, the quantitated products included the entire lane >2 nt longer than the labeled primer to exclude products resulting from NTA. Background was subtracted by using the same-sized rectangles on a portion of the screen that did not correspond to a gel lane. Fractions of product were plotted versus time and fit by a single-exponential function. For analyses of concentration dependence, rate constants were plotted against the concentration of the species being varied (RNA acceptor or dNTP in the case of NTA assays) and fit by a hyperbolic equation to obtain values of the maximal rate constant and half-maximal concentration of the varied species. Data were fitted using Prism 8 (GraphPad). Unless otherwise indicated, the reported uncertainty values reflect the standard error obtained from these fits.

### RNA-seq analysis of multiple template switches

TGIRT-seq libraries were prepared as described (34, 35) by using WT and mutant F143A GsI-IIC RT using an R2 RNA/R2R starter duplex with a 1-nt 3’-DNA overhang that is composed of equimolar mix of all four nucleotides (denoted N), and a longer 24-nt acceptor RNA, whose last three nucleotides were an equimolar mix of A, C, G, and U residues (Table S1). The initial template-switching reactions for addition of the R2R adapter to the 5′ end of the cDNA were done as described above with 200 nM WT or mutant GsI-IIC RT, 20 nM unlabeled starter duplex, and 100 nM acceptor RNA for 15 min at 60 °C. After terminating the reactions with NaOH and neutralizing with HCl as described above for template-switching reactions, the volume was raised to 100 μl with H_2_O, and cDNA products containing the R2R adapter attached to their 5′ end were cleaned-up by using a MinElute column (Qiagen) to remove unused primer. A 5′-adenylated R1R adapter was then ligated to the 3′ end of the cDNA using Thermostable 5′ App DNA/RNA Ligase (New England Biolabs) for 1 h at 65 °C. After another MinElute clean up, the entirety of the eluent was used for a PCR with an initial 5 s denaturation step at 98 °C followed by 12-cycles of 5 s denaturation at 98 °C, 10 s annealing at 60 °C, and a 10 s extension at 72 °C using Phusion polymerase (Thermo Fisher Scientific). The resulting TGIRT-seq libraries were then cleaned up by using 1.4x Ampure XP beads to remove residual primers, primer dimers, salts, and enzymes. The quality of the libraries was assessed by using a 2100 Bioanalyzer Instrument with a High Sensitivity DNA chip (Agilent).

The libraries were sequenced on an Illumina MiSeq v2 instrument to obtain 1–2 million 150-nt paired-end reads. Read 1 was used for analysis. Adaptor sequences were trimmed from read 1 using cutadapt v2.10 and then processed by using a customized R script in which each read was searched for the template sequence 5’-GCCGCTTCAGAGAGAAATCGC (3’ NNN not included) allowing up to a 2-nt deletion or insertion, 3 mismatches, and a total of 4 changes (insertions, deletions, or mismatches). Error rate were calculated for both the complete 21-nt template sequence and for a 13-nt middle (core) sequence 5’-CTTCAGAGAGAAA to avoid errors near template-switching junctions, such as those due to mismatched base pairs, extra NTAs, or heterogeneity in the length of the synthesized oligonucleotides. Sequences of the 3 randomized nucleotides between two matched template sequences were used to calculate nucleotide frequencies, and the flanking sequence upstream of the first matched template in the read (the final cDNA in template-switching product) was used for NTA analysis. The scripts used for analysis (R script) and results were deposited under https://github.com/reykeryao/Alfred.

## Data availability

*The RNA-seq data generated for this article was uploaded to BioProject ID:PRJNA723603. The structural data generated for this paper was uploaded to the Protein Data Bank with PDB ID 7K9Y. All remaining data are contained within the article*.

## References

1. Coffin, J. M. (1979) Structure, replication, and recombination of retrovirus genomes: some unifying hypotheses. J Gen Virol 42, 1–26

2. Lambowitz, A. M., and Belfort, M. (2015) Mobile Bacterial Group II Introns at the Crux of Eukaryotic Evolution. Microbiol Spectr 3, MDNA3-0050-2014

3. Shih, C., Yang, C. C., Choijilsuren, G., Chang, C. H., and Liou, A. T. (2018) Hepatitis B Virus. Trends Microbiol 26, 386–387

4. Martin-Alonso, S., Frutos-Beltran, E., and Menendez-Arias, L. (2020) Reverse Transcriptase: From Transcriptomics to Genome Editing. Trends Biotechnol

5. Inouye, S., Hsu, M. Y., Eagle, S., and Inouye, M. (1989) Reverse transcriptase associated with the biosynthesis of the branched RNA-linked msDNA in Myxococcus xanthus. Cell 56, 709–717

6. Matsuura, M., Saldanha, R., Ma, H., Wank, H., Yang, J., Mohr, G., Cavanagh, S., Dunny, G. M., Belfort, M., and Lambowitz, A. M. (1997) A bacterial group II intron encoding reverse transcriptase, maturase, and DNA endonuclease activities: biochemical demonstration of maturase activity and insertion of new genetic information within the intron. Genes Dev 11, 2910–2924

7. Kojima, K. K., and Kanehisa, M. (2008) Systematic survey for novel types of prokaryotic retroelements based on gene neighborhood and protein architecture. Mol Biol Evol 25, 1395–1404

8. Zimmerly, S., and Wu, L. (2015) An Unexplored Diversity of Reverse Transcriptases in Bacteria. Microbiol Spectr 3, MDNA3-0058-2014

9. Mestre, M. R., Gonzalez-Delgado, A., Gutierrez-Rus, L. I., Martinez-Abarca, F., and Toro, N. (2020) Systematic prediction of genes functionally associated with bacterial retrons and classification of the encoded tripartite systems. Nucleic Acids Res 48, 12632–12647

10. Liu, M., Deora, R., Doulatov, S. R., Gingery, M., Eiserling, F. A., Preston, A., Maskell, D. J., Simons, R. W., Cotter, P. A., Parkhill, J., and Miller, J. F. (2002) Reverse transcriptase-mediated tropism switching in Bordetella bacteriophage. Science 295, 2091–2094

11. Silas, S., Mohr, G., Sidote, D. J., Markham, L. M., Sanchez-Amat, A., Bhaya, D., Lambowitz, A. M., and Fire, A. Z. (2016) Direct CRISPR spacer acquisition from RNA by a natural reverse transcriptase-Cas1 fusion protein. Science 351, aad4234

12. 12. Millman, A., Bernheim, A., Stokar-Avihail, A., Fedorenko, T., Voichek, M., Leavitt, A., Oppenheimer-Shaanan, Y., and Sorek, R. (2020) Bacterial Retrons Function In Anti-Phage Defense. Cell 183, 1551–1561 e1512

13. Gao, L., Altae-Tran, H., Bohning, F., Makarova, K. S., Segel, M., Schmid-Burgk, J. L., Koob, J., Wolf, Y. I., Koonin, E. V., and Zhang, F. (2020) Diverse enzymatic activities mediate antiviral immunity in prokaryotes. Science 369, 1077–1084

14. Zhu, Y. Y., Machleder, E. M., Chenchik, A., Li, R., and Siebert, P. D. (2001) Reverse transcriptase template switching: a SMART approach for full-length cDNA library construction. Biotechniques 30, 892–897

15. Picelli, S., Bjorklund, A. K., Faridani, O. R., Sagasser, S., Winberg, G., and Sandberg, R. (2013) Smart-seq2 for sensitive full-length transcriptome profiling in single cells. Nat Methods 10, 1096–1098

16. Wulf, M. G., Maguire, S., Humbert, P., Dai, N., Bei, Y., Nichols, N. M., Correa, I. R., Jr., and Guan, S. (2019) Non-templated addition and template switching by Moloney murine leukemia virus (MMLV)-based reverse transcriptases co-occur and compete with each other. J Biol Chem 294, 18220–18231

17. Golinelli, M. P., and Hughes, S. H. (2002) Nontemplated base addition by HIV-1 RT can induce nonspecific strand transfer in vitro. Virology 294, 122–134

18. Basu, V. P., Song, M., Gao, L., Rigby, S. T., Hanson, M. N., and Bambara, R. A. (2008) Strand transfer events during HIV-1 reverse transcription. Virus Res 134, 19–38

19. Oz-Gleenberg, I., Herschhorn, A., and Hizi, A. (2011) Reverse transcriptases can clamp together nucleic acids strands with two complementary bases at their 3’-termini for initiating DNA synthesis. Nucleic Acids Res 39, 1042–1053

20. Oz-Gleenberg, I., Herzig, E., Voronin, N., and Hizi, A. (2012) Substrate variations that affect the nucleic acid clamp activity of reverse transcriptases. FEBS J 279, 1894–1903

21. Xiong, Y., and Eickbush, T. H. (1990) Origin and evolution of retroelements based upon their reverse transcriptase sequences. EMBO J 9, 3353–3362

22. Blocker, F. J., Mohr, G., Conlan, L. H., Qi, L., Belfort, M., and Lambowitz, A. M. (2005) Domain structure and three-dimensional model of a group II intron-encoded reverse transcriptase. RNA 11, 14–28

23. Mohr, S., Ghanem, E., Smith, W., Sheeter, D., Qin, Y., King, O., Polioudakis, D., Iyer, V. R., Hunicke-Smith, S., Swamy, S., Kuersten, S., and Lambowitz, A. M. (2013) Thermostable group II intron reverse transcriptase fusion proteins and their use in cDNA synthesis and next-generation RNA sequencing. RNA 19, 958–970

24. Stamos, J. L., Lentzsch, A. M., and Lambowitz, A. M. (2017) Structure of a Thermostable Group II Intron Reverse Transcriptase with Template-Primer and Its Functional and Evolutionary Implications. Mol Cell 68, 926–939 e924

25. Hu, W. S., and Hughes, S. H. (2012) HIV-1 reverse transcription. Cold Spring Harb Perspect Med 2

26. Wu, J., Liu, W., and Gong, P. (2015) A Structural Overview of RNA-Dependent RNA Polymerases from the Flaviviridae Family. Int J Mol Sci 16, 12943–12957

27. Lentzsch, A. M., Yao, J., Russell, R., and Lambowitz, A. M. (2019) Template-switching mechanism of a group II intron-encoded reverse transcriptase and its implications for biological function and RNA-Seq. J Biol Chem 294, 19764–19784

28. Qu, G., Kaushal, P. S., Wang, J., Shigematsu, H., Piazza, C. L., Agrawal, R. K., Belfort, M., and Wang, H. W. (2016) Structure of a group II intron in complex with its reverse transcriptase. Nat Struct Mol Biol 23, 549–557

29. Zhao, C., and Pyle, A. M. (2016) Crystal structures of a group II intron maturase reveal a missing link in spliceosome evolution. Nat Struct Mol Biol 23, 558–565

30. Haack, D. B., Yan, X., Zhang, C., Hingey, J., Lyumkis, D., Baker, T. S., and Toor, N. (2019) Cryo-EM Structures of a Group II Intron Reverse Splicing into DNA. Cell 178, 612–623 e612

31. Sarafianos, S. G., Das, K., Ding, J., Boyer, P. L., Hughes, S. H., and Arnold, E. (1999) Touching the heart of HIV-1 drug resistance: the fingers close down on the dNTP at the polymerase active site. Chem Biol 6, R137–146

32. Johnson, K. A. (2010) The kinetic and chemical mechanism of high-fidelity DNA polymerases. Biochim Biophys Acta 1804, 1041–1048

33. Gao, G., Orlova, M., Georgiadis, M. M., Hendrickson, W. A., and Goff, S. P. (1997) Conferring RNA polymerase activity to a DNA polymerase: a single residue in reverse transcriptase controls substrate selection. Proc Natl Acad Sci U S A 94, 407–411

34. Nottingham, R. M., Wu, D. C., Qin, Y., Yao, J., Hunicke-Smith, S., and Lambowitz, A. M. (2016) RNA-seq of human reference RNA samples using a thermostable group II intron reverse transcriptase. RNA 22, 597–613

35. Qin, Y., Yao, J., Wu, D. C., Nottingham, R. M., Mohr, S., Hunicke-Smith, S., and Lambowitz, A. M. (2016) High-throughput sequencing of human plasma RNA by using thermostable group II intron reverse transcriptases. RNA 22, 111–128

36. Xu, H., Yao, J., Wu, D. C., and Lambowitz, A. M. (2019) Improved TGIRT-seq methods for comprehensive transcriptome profiling with decreased adapter dimer formation and bias correction. Sci Rep 9, 7953

37. Yao, J., Wu, D. C., Nottingham, R. M., and Lambowitz, A. M. (2020) Identification of protein-protected mRNA fragments and structured excised intron RNAs in human plasma by TGIRT-seq peak calling. Elife 9

38. Boivin, V., Deschamps-Francoeur, G., Couture, S., Nottingham, R. M., Bouchard-Bourelle, P., Lambowitz, A. M., Scott, M. S., and Abou-Elela, S. (2018) Simultaneous sequencing of coding and noncoding RNA reveals a human transcriptome dominated by a small number of highly expressed noncoding genes. RNA 24, 950–965

39. Das, K., Martinez, S. E., Bandwar, R. P., and Arnold, E. (2014) Structures of HIV-1 RT-RNA/DNA ternary complexes with dATP and nevirapine reveal conformational flexibility of RNA/DNA: insights into requirements for RNase H cleavage. Nucleic Acids Res 42, 8125–8137

40. Nowak, E., Miller, J. T., Bona, M. K., Studnicka, J., Szczepanowski, R. H., Jurkowski, J., Le Grice, S. F., and Nowotny, M. (2014) Ty3 reverse transcriptase complexed with an RNA-DNA hybrid shows structural and functional asymmetry. Nat Struct Mol Biol 21, 389–396

41. Zhao, C., Liu, F., and Pyle, A. M. (2018) An ultraprocessive, accurate reverse transcriptase encoded by a metazoan group II intron. RNA 24, 183–195

42. Stamos, J. L., Lentzsch, A. M., Park, S. K., Mohr, G., Lambowitz, A. M. (2020) Non-ltr-retroelement reverse transcriptase and uses thereof. PCT/US2018/05414

43. Jamburuthugoda, V. K., and Eickbush, T. H. (2014) Identification of RNA binding motifs in the R2 retrotransposon-encoded reverse transcriptase. Nucleic Acids Res 42, 8405–8415

44. Bibillo, A., and Eickbush, T. H. (2002) The reverse transcriptase of the R2 non-LTR retrotransposon: continuous synthesis of cDNA on non-continuous RNA templates. J Mol Biol 316, 459–473

45. Bibillo, A., and Eickbush, T. H. (2004) End-to-end template jumping by the reverse transcriptase encoded by the R2 retrotransposon. J Biol Chem 279, 14945–14953

46. Kuiper, M. T., and Lambowitz, A. M. (1988) A novel reverse transcriptase activity associated with mitochondrial plasmids of Neurospora. Cell 55, 693–704

47. Chen, B., and Lambowitz, A. M. (1997) De novo and DNA primer-mediated initiation of cDNA synthesis by the mauriceville retroplasmid reverse transcriptase involve recognition of a 3’ CCA sequence. J Mol Biol 271, 311–332

48. Baidyaroy, D., Hausner, G., and Bertrand, H. (2012) In vivo conformation and replication intermediates of circular mitochondrial plasmids in Neurospora and Cryphonectria parasitica. Fungal Biol 116, 919–931

49. Akins, R. A., Kelley, R. L., and Lambowitz, A. M. (1986) Mitochondrial plasmids of Neurospora: integration into mitochondrial DNA and evidence for reverse transcription in mitochondria. Cell 47, 505–516

50. Lai, M. M. (1992) RNA recombination in animal and plant viruses. Microbiol Rev 56, 61–79

51. Cheng, C. P., and Nagy, P. D. (2003) Mechanism of RNA recombination in carmo- and tombusviruses: evidence for template switching by the RNA-dependent RNA polymerase in vitro. J Virol 77, 12033–12047

52. Cheng, C. P., Panavas, T., Luo, G., and Nagy, P. D. (2005) Heterologous RNA replication enhancer stimulates in vitro RNA synthesis and template-switching by the carmovirus, but not by the tombusvirus, RNA-dependent RNA polymerase: implication for modular evolution of RNA viruses. Virology 341, 107–121

53. Arnold, J. J., and Cameron, C. E. (1999) Poliovirus RNA-dependent RNA polymerase (3Dpol) is sufficient for template switching in vitro. J Biol Chem 274, 2706–2716

54. Ranjith-Kumar, C. T., Sarisky, R. T., Gutshall, L., Thomson, M., and Kao, C. C. (2004) De novo initiation pocket mutations have multiple effects on hepatitis C virus RNA-dependent RNA polymerase activities. J Virol 78, 12207–12217

55. Wang, D., Jiang, A., Feng, J., Li, G., Guo, D., Sajid, M., Wu, K., Zhang, Q., Ponty, Y., Will, S., Liu, F., Yu, X., Li, S., Liu, Q., Yang, X. L., Guo, M., Li, X., Chen, M., Shi, Z. L., Lan, K., Chen, Y., and Zhou, Y. (2021) The SARS-CoV-2 subgenome landscape and its novel regulatory features. Mol Cell 81, 1–13

56. Appleby, T. C., Perry, J. K., Murakami, E., Barauskas, O., Feng, J., Cho, A., Fox, D., 3rd, Wetmore, D. R., McGrath, M. E., Ray, A. S., Sofia, M. J., Swaminathan, S., and Edwards, T. E. (2015) Structural basis for RNA replication by the hepatitis C virus polymerase. Science 347, 771–775

57. Wang, Q., Wu, J., Wang, H., Gao, Y., Liu, Q., Mu, A., Ji, W., Yan, L., Zhu, Y., Zhu, C., Fang, X., Yang, X., Huang, Y., Gao, H., Liu, F., Ge, J., Sun, Q., Yang, X., Xu, W., Liu, Z., Yang, H., Lou, Z., Jiang, B., Guddat, L. W., Gong, P., and Rao, Z. (2020) Structural Basis for RNA Replication by the SARS-CoV-2 Polymerase. Cell 182, 417–428 e413

58. Ferrer-Orta, C., Arias, A., Perez-Luque, R., Escarmis, C., Domingo, E., and Verdaguer, N. (2007) Sequential structures provide insights into the fidelity of RNA replication. Proc Natl Acad Sci U S A 104, 9463–9468

59. Shaik, M. M., Bhattacharjee, N., Feliks, M., Ng, K. K., and Field, M. J. (2017) Norovirus RNA-dependent RNA polymerase: A computational study of metal-binding preferences. Proteins 85, 1435–1445

60. Choi, K. H., Groarke, J. M., Young, D. C., Kuhn, R. J., Smith, J. L., Pevear, D. C., and Rossmann, M. G. (2004) The structure of the RNA-dependent RNA polymerase from bovine viral diarrhea virus establishes the role of GTP in de novo initiation. Proc Natl Acad Sci U S A 101, 4425–4430

61. Kabsch, W. (2010) Xds. Acta Crystallogr D Biol Crystallogr 66, 125–132

62. Evans, P. R., and Murshudov, G. N. (2013) How good are my data and what is the resolution? Acta Crystallographica Section D 69, 1204–1214

63. Murshudov, G. N., Skubak, P., Lebedev, A. A., Pannu, N. S., Steiner, R. A., Nicholls, R. A., Winn, M. D., Long, F., and Vagin, A. A. (2011) REFMAC5 for the refinement of macromolecular crystal structures. Acta Crystallogr D Biol Crystallogr 67, 355–367

64. Emsley, P., Lohkamp, B., Scott, W. G., and Cowtan, K. (2010) Features and development of Coot. Acta Crystallogr D Biol Crystallogr 66, 486–501

