## Supplemental information for "Structural basis for template switching by a group II intron-encoded non-LTR-retroelement reverse transcriptase"

**Table S1. DNA and RNA oligonucleotides used in this study**

**Crystallography**

| **Donor duplex** | |
|---|---|
| 053 (DNA primer) | 5'-CTCCAGGCAAddC |
| r001-4 (donor RNA template) | 5'-UUGCCUGGAG |

| **Acceptor RNA** | |
|---|---|
| rUUUUG | 5'-UUUUG |

**Biochemical assays**

| **Starter duplex** | |
|---|---|
| R2R-G (DNA primer) | 5'-GTGACTGGAGTTCAGACGTGTGCTCTTCCGATCTG |
| R2R-Blunt (DNA primer) | 5'-GTGACTGGAGTTCAGACGTGTGCTCTTCCGATCT |
| R2 (donor RNA template) | 5'-AGAUCGGAAGAGCACACGUCUGAACUCCAGUCAC |

| **Acceptor RNA** | |
|---|---|
| 21-nt acceptor RNA | 5'-GCCGCUUCAGAGAGAAAUCGC |
| 34-nt acceptor RNA | 5'-CAGACGAGCUCAUGCCGCUUCAGAGAGAAAUCGC |

| | |
|---|---|
| Primer extension DNA primer | 5'-TCTTCGGGGCGAAAACTCTCAAGGATCTTACCGC TGTTGAGATCCAGTTC |

**TGIRT-seq**

| **Starter duplex** | |
|---|---|
| R2R-N (DNA primer)[1] | 5'-GTGACTGGAGTTCAGACGTGTGCTCTTCCGATCTN |
| R2 (RNA template) | 5'-AGAUCGGAAGAGCACACGUCUGAACUCCAGUCAC |

| **Acceptor RNA** | |
|---|---|
| 24-nt acceptor RNA[2] | 5'-GCCGCUUCAGAGAGAAAUCGCNNN |

[1] N denotes equimolar A, C, G, and T residues obtained by hand mixing of four separate oligonucleotides.
[2] NNN denotes oligonucleotides synthesized with machine-mixed equimolar A, C, G, and U residues.

**Table S2. WT and F143A GsI-IIC RT error rates.**

**21-nt constant region (error frequency x $10^{-3}$)**

| Error type | WT (4 mM) | WT (0.4 mM) | F143A (4 mM) | F143A (0.4 mM) |
|---|---|---|---|---|
| Substitution | 6.51 | 5.18 | 4.84 | 5.12 |
| Insertion | 0.93 | 0.85 | 0.97 | 0.90 |
| Deletion | 1.45 | 1.26 | 1.23 | 1.63 |
| Overall | 8.89 | 7.29 | 7.04 | 7.64 |

**13-nt internal region (error frequency x $10^{-3}$)**

| Error type | WT (4 mM) | WT (0.4 mM) | F143A (4 mM) | F143A (0.4 mM) |
|---|---|---|---|---|
| Substitution | 5.18 | 4.14 | 4.07 | 3.41 |
| Insertion | 0.66 | 0.64 | 0.74 | 0.69 |
| Deletion | 0.34 | 0.29 | 0.28 | 0.28 |
| Overall | 6.18 | 5.08 | 5.09 | 4.37 |

The Table shows error frequencies for TGIRT-seq of a 24-nt acceptor RNA with three N nucleotides at its 3' end using WT and mutant F143A GsI-IIC RTs with template switching and reverse transcription done at 4 mM and 0.4 mM dNTPs. The error rates were determined for the entire 21-nt constant region and a 13-nt internal region ("CTTCAGAGAGAAA") of the 24-nt acceptor.

#### Figure S1

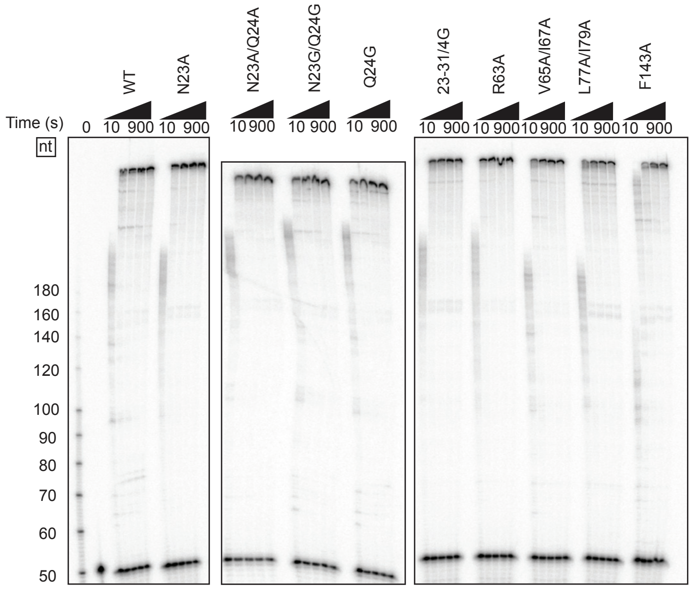


| RT | Amplitude | Rate (nt/s) |
|---|---|---|
| WT | 0.88 (0.03) | ~ 20 |
| N23A | 0.86 (0.02) | ~ 20 |
| N23AQ24A | 0.89 (0.00) | ~ 20 |
| N23GQ24G | 0.90 (0.00) | ~ 20 |
| Q24G | 0.90 (0.01) | ~ 20 |
| 23-31/4G | 0.86 (0.02) | ~ 20 |
| R63A | 0.85 (0.02) | ~ 20 |
| V65A/I67A | 0.85 (0.02) | ~ 15 |
| L77A/I79A | 0.88 (0.01) | ~ 15 |
| F143A | 0.88 (0.01) | ~ 10 |

**Figure S1. Primer-extension activity of wild-type and mutant GsI-IIC RTs in this study.** Time courses of primer-extension reactions were done with 200 nM wild-type (WT) or mutant GsI-IIC RTs, 20 nM 1.1-kb RNA template annealed to a 50-nt 5′-$^{32}$P-labeled DNA primer, and 4 mM dNTPs (an equimolar mix of 1 mM dATP, dCTP, dGTP, and dTTP) in reaction medium containing 200 mM NaCl at 60 °C. Aliquots were quenched at times ranging from 10 to 900 s, and the products were analyzed in a denaturing 6% poly-acrylamide gel. The numbers to the left of the gel indicate sizes of 5′-$^{32}$P-labeled single-stranded DNA size markers (ss20 DNA Ladder, Simplex Sciences) run in a parallel lane. The assays for each mutant were performed three times. Amplitudes were calculated as the proportion of $^{32}$P label in full-length product compared to total $^{32}$P-label in the lane, with standard deviations in parentheses. Rates in nucleotides per second were approximated by the progression of the center of the distribution at the 10 s timepoint.

### Figure S2

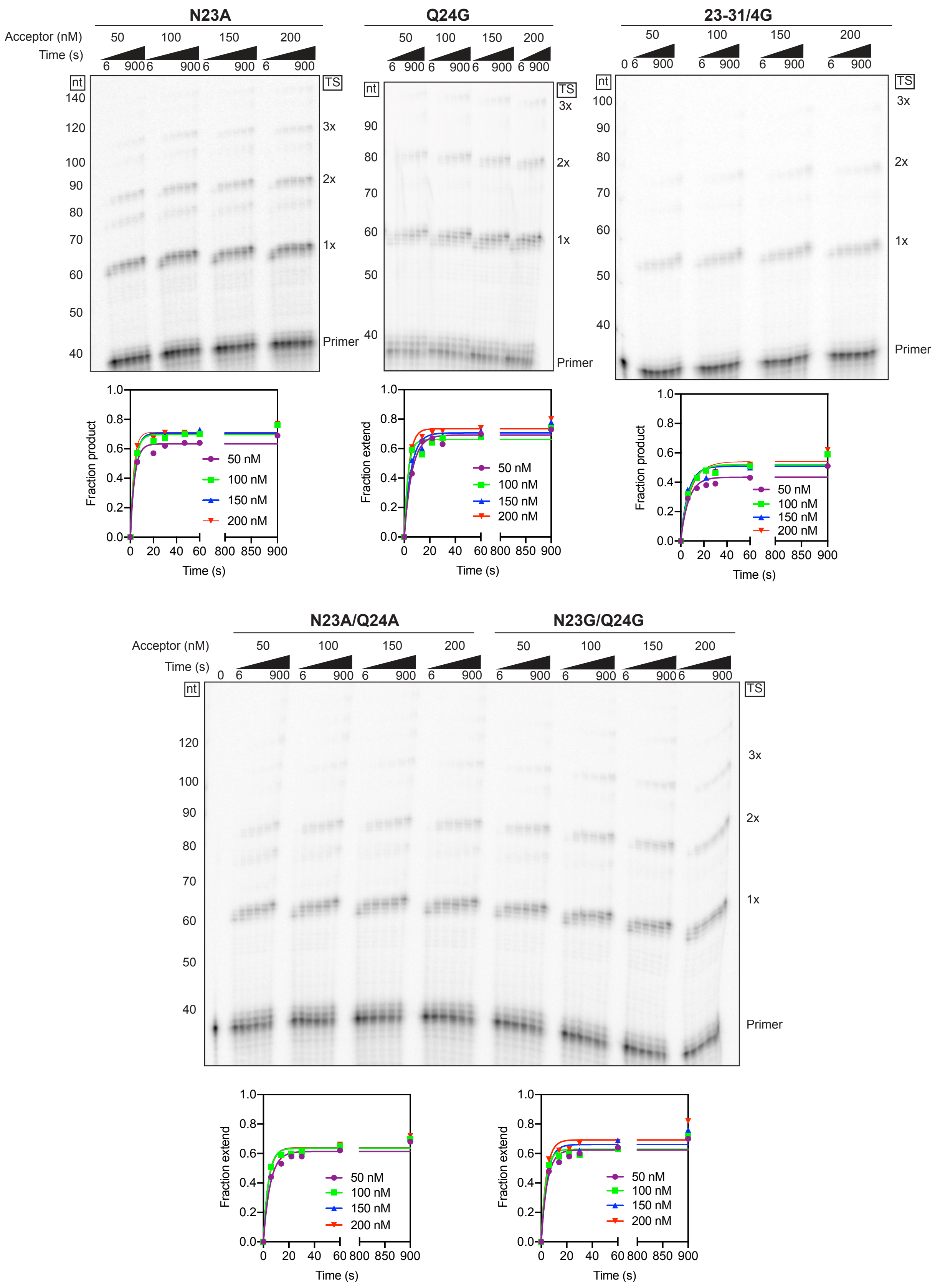

N23A
Q24G
23-31/4G
Acceptor (nM)
Time (s)
50
100
150
200
6
900
0
nt
TS
140
120
100
90
80
70
60
50
40
3x
2x
1x
Primer
1.0
0.8
0.6
0.4
0.2
0.0
Fraction product
Fraction extend
50 nM
100 nM
150 nM
200 nM
0
20
40
60
800
850
900
Time (s)
N23A/Q24A
N23G/Q24G

**Figure S2. Time courses of template switching as a function of acceptor RNA concentration used to determine kinetic parameters for GsI-IIC RT RT0 loop mutants.** The Figure shows representative gels and plots of template-switching reaction time courses done as described in Fig. 3. The names of mutants are indicated above the gel. The numbers to the left of the gels indicate the length of size markers (5′-$^{32}$P-labeled single-stranded DNA ladder; ss20 DNA Ladder, Simplex Sciences) run in a parallel lane. 1x, 2x and 3x to the right of the gel indicate the number of consecutive template switches for the product bands at these positions. The plots below each gel show time courses of production of template-switching products (*i.e.*, products >2 nt longer than the primer) at each RNA acceptor concentration, with the data fit by a single-exponential function.

#### Figure S3

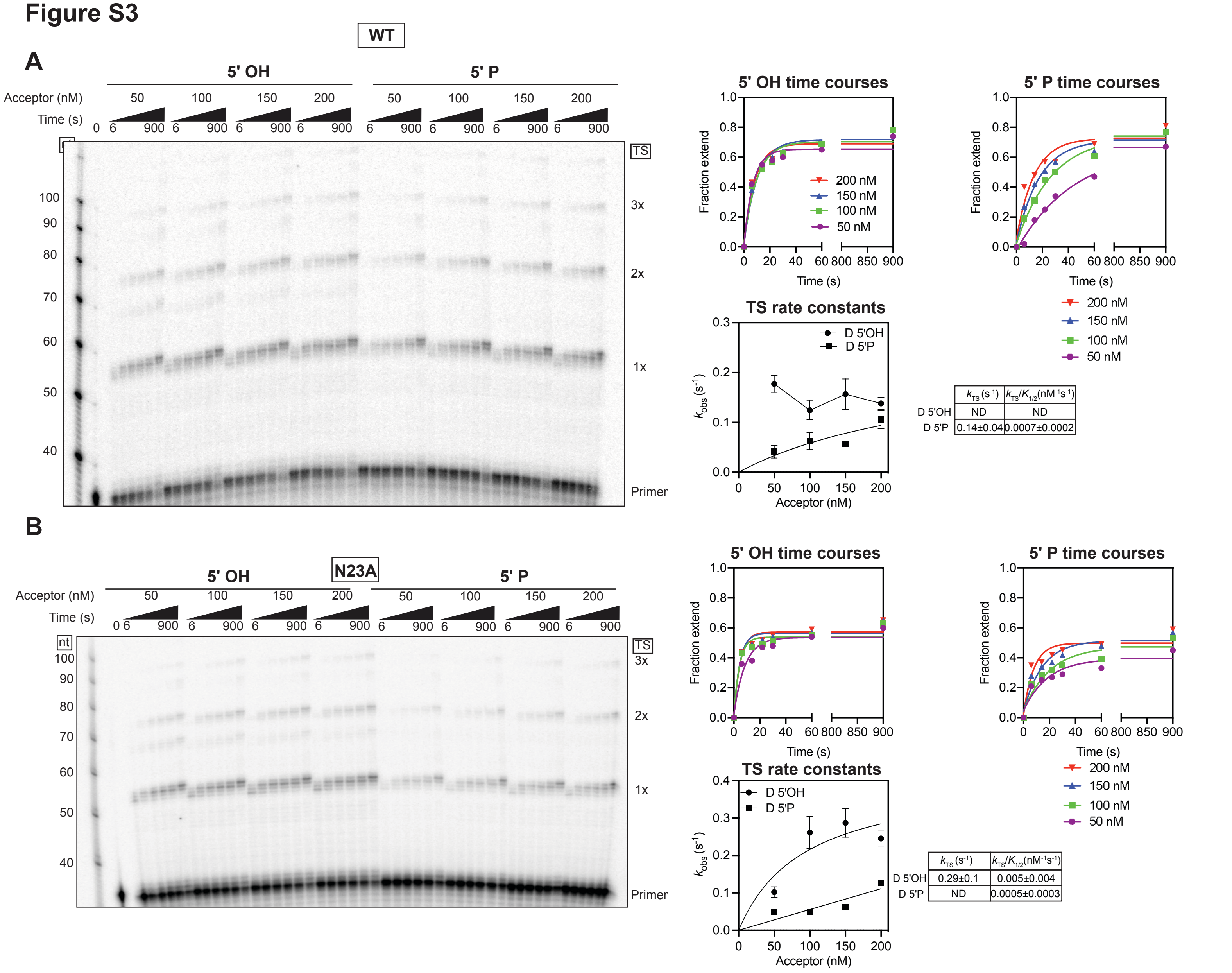


**Figure S3. Kinetic parameters for template switching of wild-type and N23A mutant GsI-IIC RTs from donor RNA templates having a 5' OH or 5' phosphate.** *A* and *B*, time courses of template switching as a function of acceptor RNA concentration for wild-type (WT) and N23A mutant GsI-IIC RTs, respectively, with donor RNA template/DNA primer duplexes having a 5' OH or 5' phosphate (5' P) RNA template. Template-switching reactions were done as described in Fig. 3, and the products were analyzed in a denaturing 6% polyacrylamide gel. The numbers to the left of the gels indicate the lengths of size markers (5′-$^{32}$P-labeled single-stranded DNA ladder; ss20 DNA Ladder, Simplex Sciences) run in the leftmost lane, and 1x, 2x and 3x to the right of the gel indicate the number of consecutive template switches for the product bands at these positions. Plots for the time courses are shown at the top right of the gel, with plots of the observed rate constant ($k_{obs}$) as a function of acceptor concentration fit by a hyperbolic function shown below. Each rate constant determination was performed two to four times.

#### Figure S4

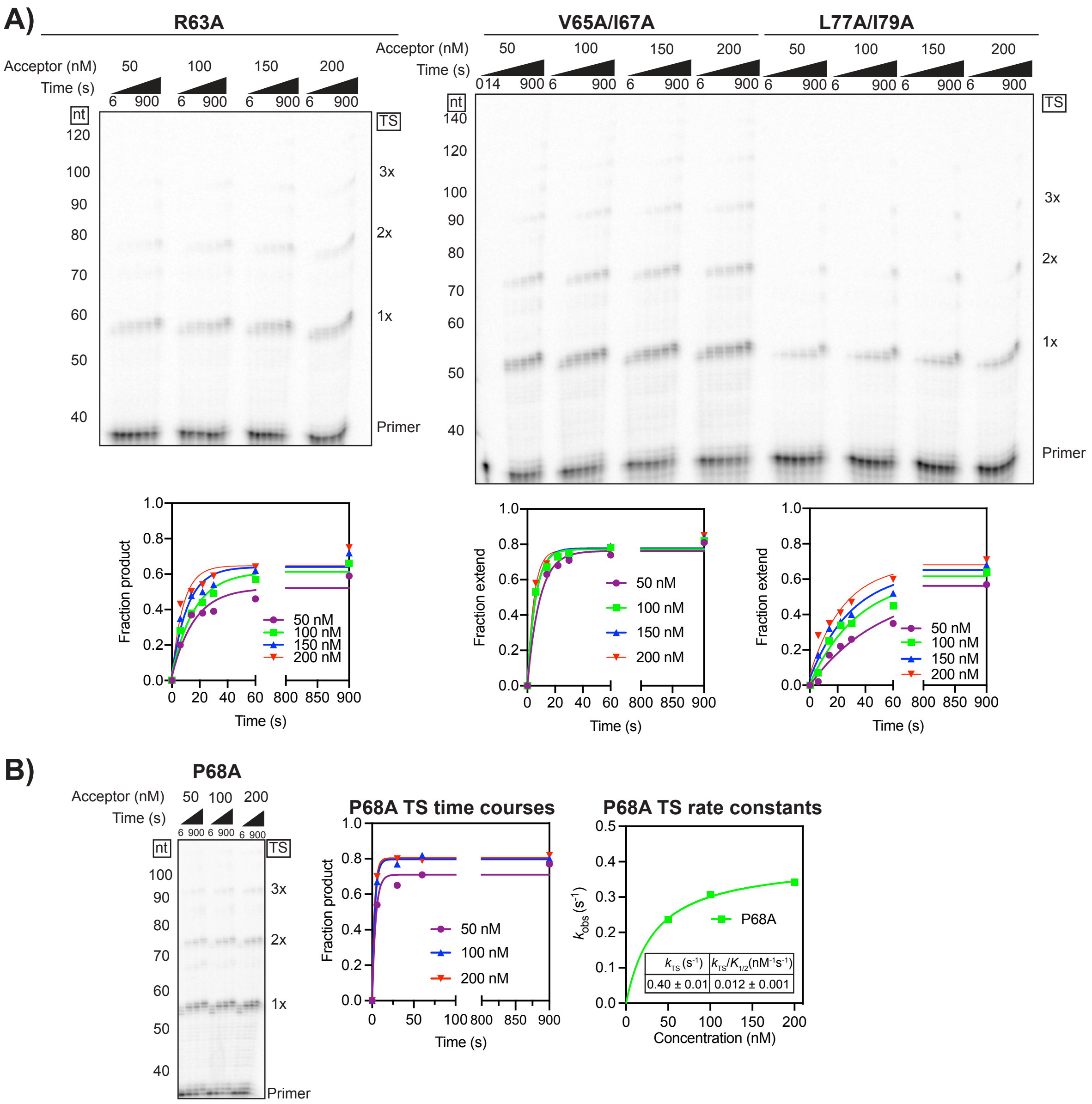


**Figure S4. Time courses of template switching as a function of acceptor RNA concentration used to determine kinetic parameters for GsI-IIC RT fingertips loop mutants.** *A*, representative gels and plots of template-switching reaction time courses for mutants R63A, V65A/I67A, and L77A/I79A. *B*, gel and plots of template-switching reaction time courses and of the observed rate constant ($k_{obs}$) as a function of acceptor concentration fit by a hyperbolic function for mutant P68A. Template-switching reactions were done as described in Fig. 3. The numbers to the left of the gels indicate length of size markers (5′-$^{32}$P-labeled singe-stranded DNA ladder; ss20 DNA Ladder, Simplex Sciences) run in a parallel lane of the same gel. 1x, 2x and 3x to the right of the gel indicate the number of consecutive template switches for the product bands at these positions.

Figure S5

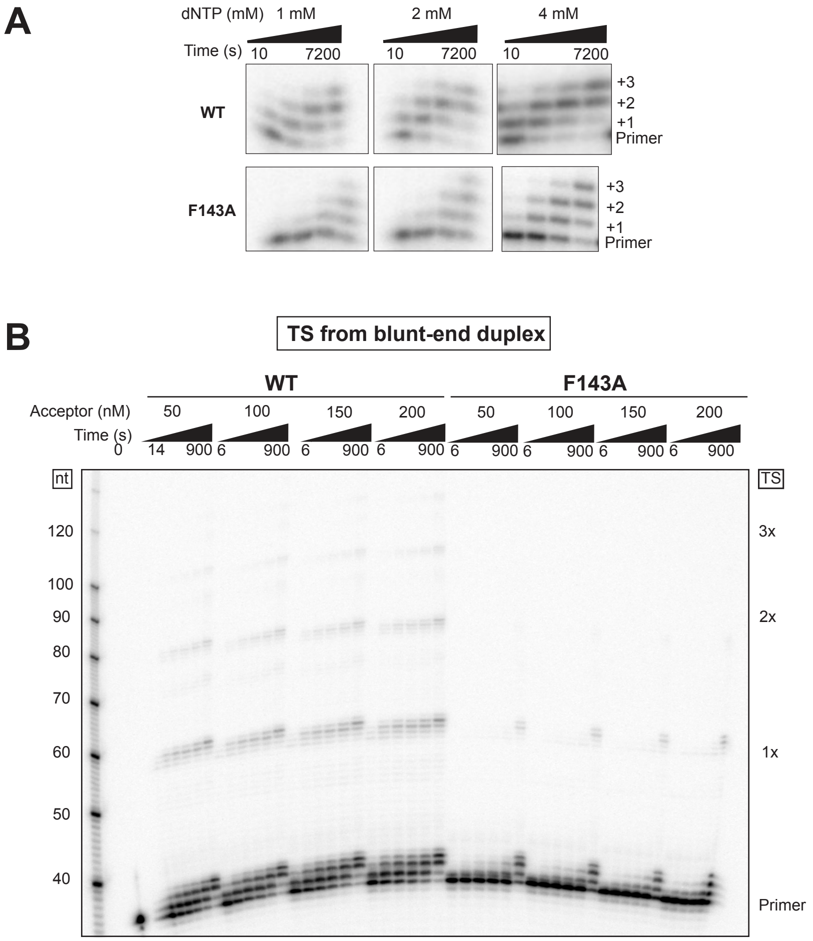


**Figure S5. Time courses for NTA and template-switching reactions from a blunt-end duplex as a function of acceptor RNA concentration for WT and F143A mutant GsI-IIC RTs.** *A*, representative gels for time courses of NTA reactions at 1, 2, and 4 mM dNTP done as described and plotted in Fig. 6*B*. +1, +2, and +3 indicate the number of added nucleotides. *B*, representative gel for time courses of template switching from a blunt-end duplex by WT and F143A mutant GsI-IIC RTs used for the plots in Fig. 6*D*. The numbers to the left of the gel indicate lengths of size markers (5′-$^{32}$P-labeled single stranded DNA ladder; ss20 DNA Ladder, Simplex Sciences) run in the left most lane. 1x, 2x and 3x to the right of the gel indicate the number of consecutive template switches for the product bands at these positions.
